# Understanding and leveraging phenotypic plasticity during metastasis formation

**DOI:** 10.1101/2022.11.07.515430

**Authors:** Saumil Shah, Lisa-Marie Philipp, Stefano Giaimo, Susanne Sebens, Arne Traulsen, Michael Raatz

**Affiliations:** Department of Evolutionary Theory, Max Planck Institute for Evolutionary Biology, Ploen, August-Thienemann-Str. 2, 24306 Ploen; Institute for Experimental Cancer Research, Kiel University and University Hospital Schleswig-Holstein, Campus Kiel, Arnold-Heller-Str. 3, Building U30, Entrance 1, 24105 Kiel

**Keywords:** mathematical oncology, phenotypic plasticity, adaptive therapy, epithelial mesenchymal transition, metastasis

## Abstract

Cancer metastasis is the process of detrimental systemic spread and the primary cause of cancer-related fatalities. Successful metastasis formation requires tumor cells to be proliferative and invasive; however, cells cannot be effective at both tasks simultaneously. Tumor cells compensate for this trade-off by changing their phenotype during metastasis formation through phenotypic plasticity. Given the changing selection pressures and competitive interactions that tumor cells face, it is poorly understood how plasticity shapes the process of metastasis formation. Here, we develop an ecology-inspired mathematical model with phenotypic plasticity and resource competition between phenotypes to address this knowledge gap. We find that phenotypically plastic tumor cell populations attain a stable phenotype equilibrium that maintains tumor cell heterogeneity. Considering treatment types inspired by chemo- and immunotherapy, we highlight that plasticity can protect tumors against interventions. Turning this strength into a weakness, we corroborate current clinical practices to use plasticity as a target for adjuvant therapy. We present a parsimonious view of tumor plasticity-driven metastasis that is quantitative and experimentally testable, and thus potentially improving the mechanistic understanding of metastasis and its treatment consequences.

## 1 Introduction

Cancer metastasis formation, or the spread and growth of tumor cells throughout the body, is the cause of more than 60% of cancer-related deaths ^1^. Despite many decades of drug development for cancer, the survival rate for patients with metastatic cancer remains low ^2,3^. The process of metastasis is a multistage process that includes local invasion by the tumor cells, invading into the circulatory system, survival in circulation, arrest at a distant tissue, getting out of the circulation, survival, adaptation and outgrowth in a new environment^4–6^. Tumor cells face selection for proliferation, locally at each site of cancer, and for invasiveness and motility during the spread between two sites. These changing selection pressures during metastasis are driven by environmental factors and interactions with different surroundings ^7,8^.

How tumor cells respond to these frequently changing environmental conditions is crucial for their persistence. Tumor cell populations possess several adaptive strategies to cope with such fluctuating environments. A subset of these strategies relies on diversity-generating mechanisms. In addition to genetic mechanisms, such as mutations, copy number alterations, and translocations, there are non-genetic mechanisms to generate diversity, such as epigenetic regulation, stochastic gene expression, and cellular differentiation hierarchies ^7,9^. All these factors are internal to the cell. In addition, the local environment can be substantial local variations in the tumor microenvironment may generate functional heterogeneity among clonal tumor cells ^7,8^.

An early review by West-Eberhard ^10^ emphasizes plasticity as a diversity-generating mechanism and its contribution to altering traits. Plasticity also affects population and eco-evolutionary dynamics ^11^.

One prominent outcome of epithelial tumor cell plasticity is the range of phenotypes and transitions on the spectrum from epithelium to mesenchyme. Epithelial cells are proliferative but cannot move, while mesenchymal cells are motile and invasive but proliferate slowly ^12^. Both proliferative and invasive phenotypes are crucial for tumor progression and metastasis. Thus, shifts in the phenotype of tumor cells by epithelial-mesenchymal plasticity are salient features of metastatic cancers^13–16^. Consequently, during the epithelial-to-mesenchymal transition, partial mesenchymal features are gained, and epithelial features are progressively lost, leading to altered surface marker expression, decreased cell-cell adhesion, and motility-facilitating cytoskeleton reorganiation ^17^. The reverse change happens during the mesenchymal-to-epithelial transition.

These transitions result in the presence of hybrid phenotypes along the continuous spectrum between epithelial and mesenchymal phenotypes. This continuous spectrum is often discretied into a number of hybrid phenotypes, with varying estimates on the number of these hybrid phenotypes ^18–22^. The plasticity between epithelial and mesenchymal phenotypes was found to be promoted by interactions with the tumor microenvironment. Among others, high levels of ransforming Growth actor *β* (TGF-*β*) are critical drivers toward more mesenchymal phenotypes ^8,17,23^. Different treatment types can select differently on this plasticity^24^. the current first-line treatment for many cancers is chemotherapy which targets cell proliferation. A more recent treatment option for some cancers is immunotherapy which comprises antibody-based or cellular therapies targeting molecules present on the cell membrane of the cancer cells.

These differential selection pressures suggest classifying treatments into growth-dependent or growth-independent treatments. As epithelial-like phenotypes proliferate faster, they are more sensitive to chemotherapy than mesenchymal phenotypes. Immunotherapy should target all phenotypes similarly, assuming that it does not target surface markers that change during epithelial-mesenchymal transition. Adjuvant therapies, such as TGF-*β* blockers, are promising candidates to modify the transition process and, thus, ultimately, the phenotypic plasticity of tumor cells ^25–27^. As TGF-*β* promotes the mesenchymal phenotype ^26,27^ and TGF-*β* blockers improve chemotherapy^25^, it is suspected that the gene regulatory network of the epithelial mesenchymal transition could be a key to control the balance of phenotypes and thus the treatment outcome ^28,29^. This gene regulatory network consists of cell fate controlling signaling pathways like TGF-*β*, WN and NO CH that recruit epithelial-mesenchymal-transition-inducing transcription factors ZEB, SNAIL, SLUG, TWIST or miR-200 ^17,29–32^.

Mathematical modeling presents a valuable approach to understanding the mechanisms and consequences of genetic and non-genetic diversity-generating mechanisms. Along this line, Zhou et al. ^33^ modeled the hierarchical cellular diversity created by differentiation and de-differentiation, with stem cells at the root of the hierarchy and specialized cells at its leaves. Gupta et al. ^34^ and Li and Thirumalai ^35^showed with experiments and mathematical modeling that phenotypically plastic breast cancers approach a phenotypic equilibrium and can maintain phenotypic heterogeneity. A mathematical model of phenotype switching between drug-sensitive and drug-resistant cells in non-small cell lung cancer ^36^ also shows a similar behavior. Such plastic mechanisms in tumor cells cause reduced efficacy of chemotherapy and acquired therapeutic resistance ^13,37,38^. In a spatial, multi-organ model, Franssen and Chaplain focus specifically on epithelial-mesenchymal transitions during metastatic spread without treatment^39^. In a recent paper, we modeled the phenotype switching between fast and slowly proliferating B cells during the relapse dynamics in acute lymphoblastic leukemia and investigated the impacts of treatment on phenotypic heterogeneity^24^.

Now, in this study we propose an unconform yet parsimonious view of the phenotype transitions without explicitly considering the microenvironment. This view considers the trait variation present in a tumor population and consequently the heterogeneous response to drug as an inherent property of the tumor. We also discuss feasible observables that could be measured for patient-derived tumors to test our hypothesis. In this study we focus on (i) how different characteristics of plasticity shape the phenotype distribution, (ii) how this distribution determines the transition dynamics between epithelial and mesenchymal phenotypes at primary and secondary tumor sites, and (iii) how these dynamics are affected by chemo- or immunotherapy, particularly applied as adjuvant therapy (after surgical reduction of the primary tumor mass), and additionally by a transition-modulating therapy. With the proposal of a practical transition modulating therapy we emphasize the phenotype transition as a viable clinical target.

## 2 Results

Using a dynamical model of the abundances, *x*_*i*_, of *N* distinct phenotypes with linearly decreasing growth rates along the epithelial-mesenchymal trait axis, *i* = 1, 2, …, *N*, we track the temporal phenotype dynamics before, during and after treatment (see Fig. 1). We implement competition between phenotypes by assuming a shared carrying capacity *K*. We include transitions to an adjacent epithelial-like phenotype *T*_*EM*_ and adjacent mesenchymal-like phenotype *T*_*ME*_. We propose that a molecule can control the balance between these transitions *T*_*EM*_ and *T*_*ME*_. The concentration of this transition-modulating molecule can be mapped to the transition bias *λ*. The relation between the growth and the speed of phenotype transition is encoded by transition speed *c*. Further, we investigate growth-dependent treatment *m*_*D*_, i.e., chemotherapy, and growth-independent treatment *m*_*I*_, i.e., immunotherapy, to explore treatments and administration sequences. For a detailed description of the modeling assumptions please see the Section Model.

**Figure 1:**
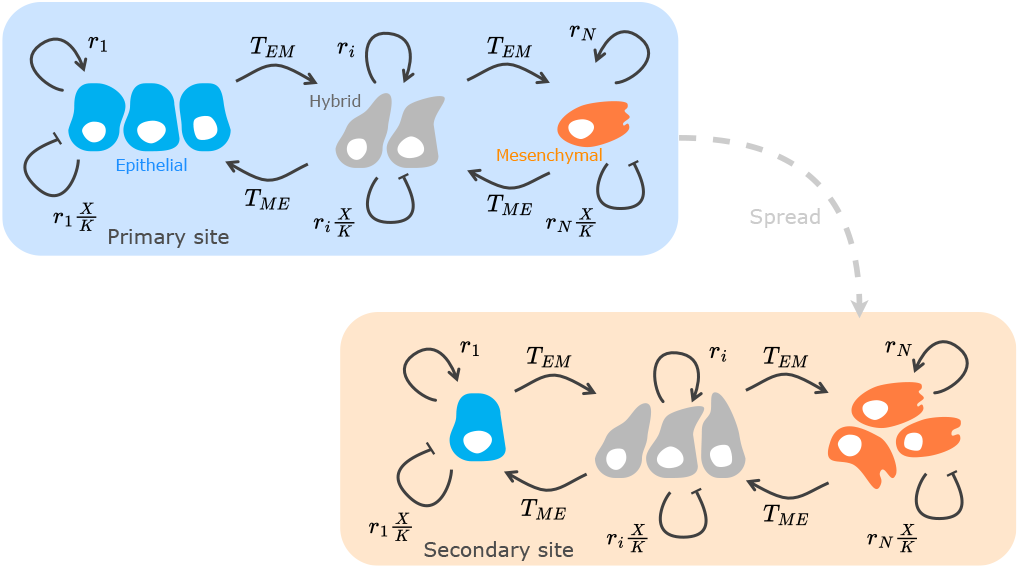
Model structure capturing phenotypic heterogeneity at the primary and secondary tumor site. We focus on the competitive growth of heterogeneous tumor cell populations at each tumor site. The dashed arrow represents spread of cancer cells between the primary and secondary sites. We do not model the spread between tumor sites explicitly, but consider the spread as translating into different initial phenotype distributions at primary and secondary sites. The different compartments in the model for each represent epithelial, hybrid, and mesenchymal phenotypes. The solid arrows indicate competitive growth and phenotype transitions. Phenotype *i* grows at rate *r*_*i*_ and transitions to the adjacent more mesenchymal-like type at rate *T*_*EM*_ and to the adjacent more epithelial-like type at rate *T*_*ME*_. Resource competition is modeled with the term 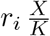 where 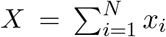 is the total population abundance, and *K* is the carrying capacity. The epithelial phenotype can only transition to the more mesenchymal-like adjacent hybrid phenotype, and the mesenchymal phenotype can only transition to the epithelial-like adjacent hybrid phenotype thus, the epithelial and mesenchymal phenotypes are the terminal phenotypes. Here, we show only one hybrid phenotype, but we also investigate the effect of a larger number of hybrid phenotypes.

### Phenotype transitions generate and maintain heterogeneity

Assuming that cells can transition into adjacent epithelial or mesenchymal phenotypes generates a phenotype distribution of abundances along the epithelial-mesenchymal trait axis. Without treatment, *m*_*D*_ = *m*_*I*_ = 0, the model (see Section Model) has two equilibria. One equilibrium is the state where the tumor vanishes, *x*_*i*_ = 0 for *i* = 1, 2, …, *N*. This equilibrium is unstable, implying that a tumor can always progress and increase in abundance without treatment. The second equilibrium is the state described by

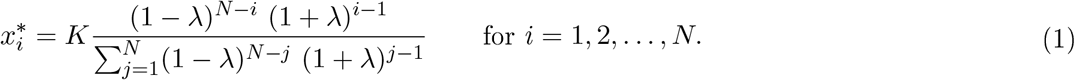

This equilibrium is stable and describes the coexistence of all phenotypes (see supplementary text for a proof of global stability). This stability implies that phenotypic heterogeneity is generated and maintained in tumor populations featuring phenotypic transitions. Patient-derived breast cancer cell populations exhibit similar behavior _*in-vitro*_ ^34,35^.

The sum of all phenotype abundances at the coexistence equilibrium is *K*. The stable distribution of phenotypes at the coexistence equilibrium depends on the transition bias *λ* and the number of phenotypes *N*. Notably, the stable phenotype distribution does not depend on the growth rate. Thus, only the transitions and not the growth dynamics decide the phenotype abundances at equilibrium. Additionally, this growth independence shows that the particular choice of the decline of growth rates from epithelial to mesenchymal phenotypes does not affect the equilibrium phenotype distribution.

### The transition bias controls the stable phenotype distribution

When cells change to a more epithelial or mesenchymal phenotype with the same probability, i.e., when there is no transition bias (*λ* = 0), all phenotypes are at equal abundance and the stable distribution is a uniform distribution (Fig. 2). The mesenchymal phenotype (M) is the most abundant when there is a transition bias towards the mesenchymal phenotypes (*λ >* 0). On the other hand, for a transition bias towards the epithelial phenotypes (*λ <* 0), the epithelial phenotype (E) is the most abundant. The phenotypic heterogeneity of the tumor population is highest when there is no transition bias and *λ* = 0 (Fig. S1). A high variance corresponds to presence of a lot of different phenotypes reducing the efficacy of standard cancer treatments. The heterogeneity decreases and vanishes at the bias extremes *λ* → ±1 where transitions become unidirectional.

**Figure 2:**
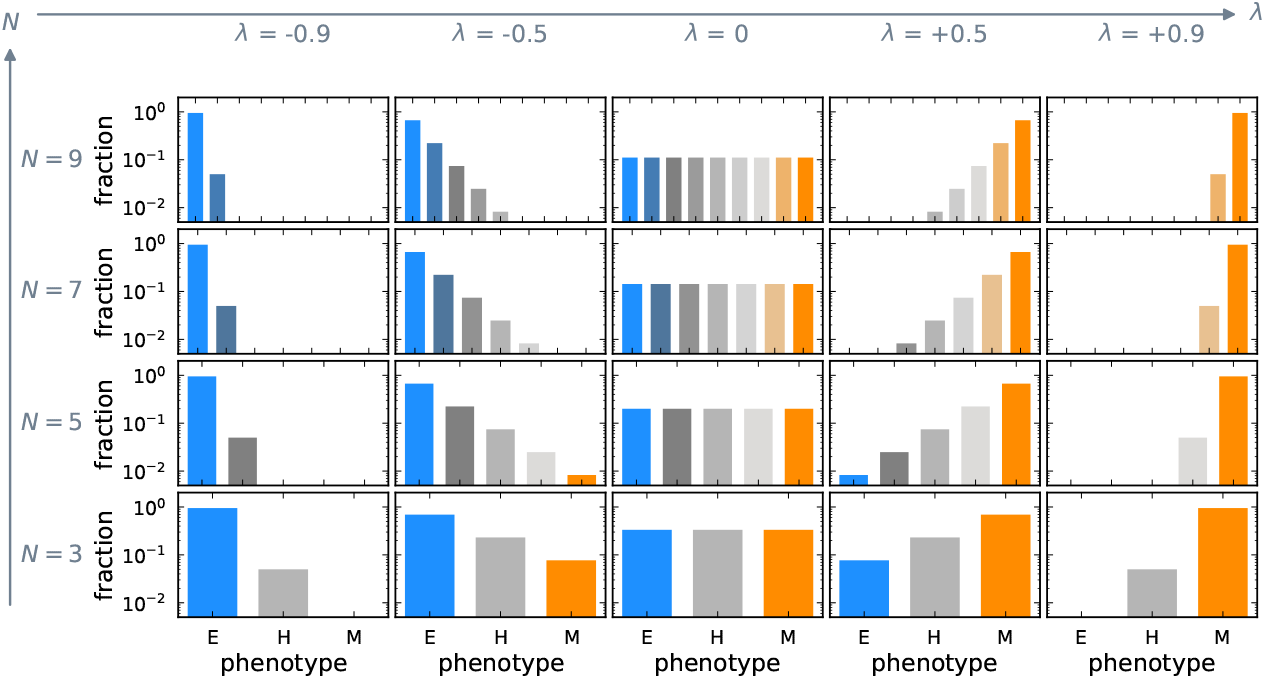
Phenotype transitions create a stable phenotype distribution. Each panel shows the distribution of phenotypes (Eq. (1)) relative to the carrying capacity *K* for a fixed value of the transition bias *λ* and the number of phenotypes *N*. The stable phenotype distribution changes with the transition bias *λ* but remains qualitatively unaffected by changing the number of phenotypes *N*. The stable distribution is uniform when there is no transition bias to either epithelial or mesenchymal-like phenotypes, i.e., *λ* = 0. *λ <* 0 depicts a transition bias towards epithelial-like phenotypes and leads to a relative increase in epithelial cells. Conversely, *λ >* 0 results in a transition bias towards mesenchymal-like phenotypes and causes a relative increase in mesenchymal cells.

The presence of phenotypes far off the average population phenotype with very low abundances is captured by the third central moment of the phenotype distribution. These phenotypes are essential for residual disease in some cancers, and thus an important potential treatment target. We find that the third central moment is an odd function of the transition bias, vanishing at *λ* = 0 and at high transition bias (Fig. S1). The number of phenotypes *N* quantitatively affects the moments of the stable phenotype distribution but does not affect their shapes qualitatively (Fig. S1). We will thus fix *N* = 3 phenotypes in the remainder of the study for illustration.

### The transition speed determines how fast the stable phenotype distribution is approached

While the transition bias *λ* determines the stable phenotype distribution, the transition speed *c* affects how fast the stable phenotype distribution is approached. The model consists of two processes logistic growth and transitions, where the parameter *c* scales the transition propensity relative to the growth rate. When transitions are faster than growth, *c >* 1, the population first reaches the stable phenotype distribution and then approaches the carrying capacity. When transitions are slower than growth, *c <* 1, the population first reaches the carrying capacity and then equilibrates to the stable phenotype distribution (Fig. 3).

**Figure 3:**
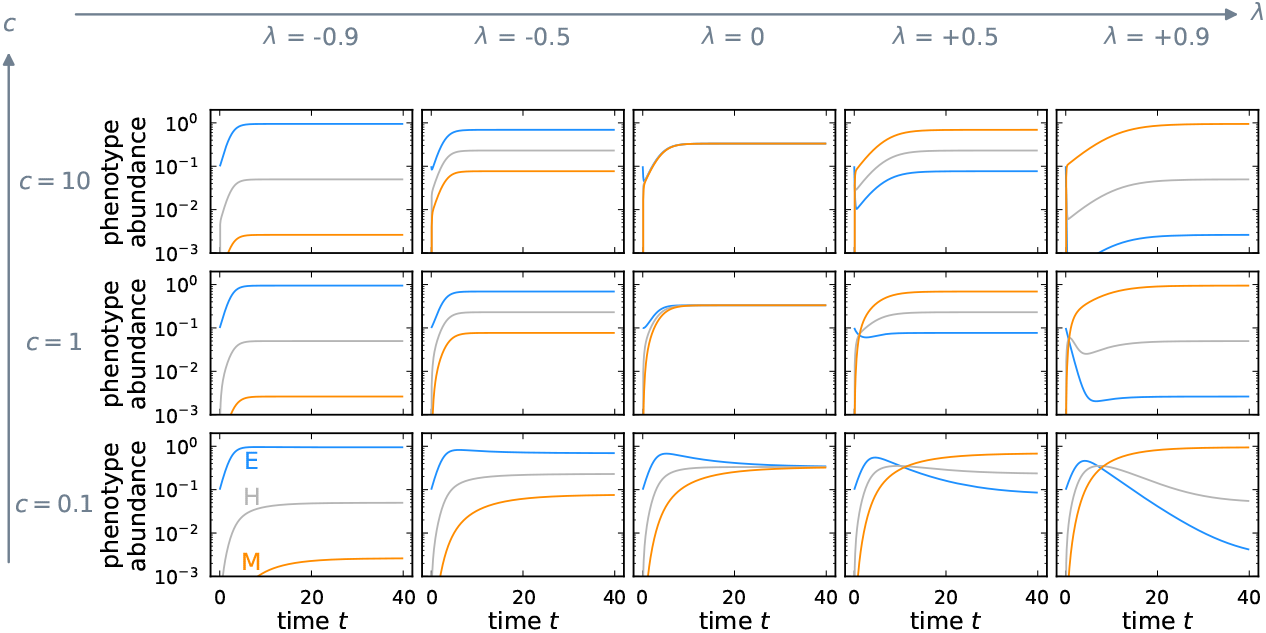
The rate of approach to stable phenotype distribution is determined by the transition speed. The panels show the approach to the stable phenotype distribution from the same initial condition, 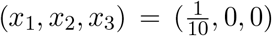, for different combinations of transition speed *c* and transition bias *λ*. The transition speed *c* sets the pace of the transition dynamics relative to the growth dynamics. For any transition bias *λ*, the time to reach the stable phenotype distribution decreases as the transition speed increases.

### The initial phenotype distribution determines the direction of phenotype shift

The stability of the coexistence equilibrium induces phenotype shifts similar to epithelial-mesenchymal and mesenchymal-epithelial transitions. The primary site of a tumor at an early stage is composed mainly of proliferative epithelial cells. Thus, at the primary site, the stable phenotype distribution is approached by a relative increase in the abundance of mesenchymal-like cells (Fig. 4, first column). The secondary tumor site is initially founded by mesenchymal-like cells due to the selection pressure for invasive properties during dissemination. Therefore, conversely to the primary site, the stable equilibrium is approached by a relative increase of epithelial-like cells at the secondary site (Fig. 4, second column) during metastasis formation.

**Figure 4:**
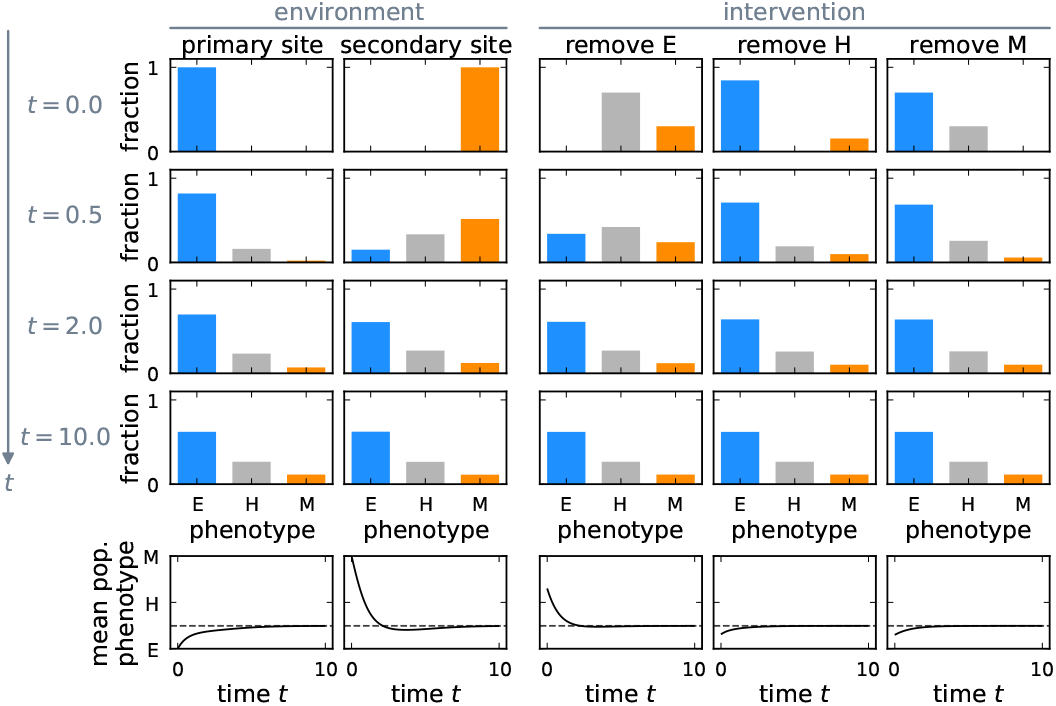
Plasticity-driven approach to stable phenotype distribution from different initial conditions. Depending on the initial condition, the approach to the stable phenotype distribution proceeds along different paths (top four rows, the vertical axis represents time). The mean population phenotype over time is shown in the last row with its asymptote (dashed line). The first column shows the phenotype dynamics of a plastic tumor at the primary site, where it originates only from epithelial cells. The second column represents the growth of a metastasis after mainly mesenchymal cells have arrived at the secondary site. The last three columns show the dynamics after a hypothetical intervention that removes all epithelial, all hybrid, or all mesenchymal cells. In all cases, the tumor approaches the stable phenotype distribution however, different initial conditions lead to shifting the average phenotype in different directions.

After a perturbation from the coexistence equilibrium, e.g., by treatment, the stable phenotype distribution is restored, compensating the effect of the perturbation (Fig. 4, columns three to five). Hence, the phenotypically plastic tumor cell population can maintain heterogeneity. Our model thus exhibits site- and treatment-specific phenotype shifts that result only from the selection during dissemination or treatment. These shifts do not require specific microenvironmental factors, although the microenvironment is a crucial component contributing to the initial conditions that determine the direction of the shift.

### Treatment type affects the phenotype distribution of the tumor

Next, we tested how the phenotype distribution changes during and after growth-dependent and growth-independent treatment (Fig. 5). Growth-dependent treatment targets proliferating cells, thus epithelial cells experience higher treatment-induced mortality (Fig. 5**a**), and the mean of the distribution shifts towards the mesenchymal phenotype (Fig. 5**c**). Growth-independent treatment targets all phenotypes equally, not changing the phenotype distribution directly. The room for growth is filled by epithelial-like cells as they grow fast (Fig. 5**b**) during treatment as the tumor is not at carrying capacity. Thus, during growth-independent treatment, the phenotype average shifts towards the epithelial phenotype, especially, if transitions are slow compared to growth dynamics (*c <* 1, Fig. 5**d**). After treatment, the phenotype composition returns to the stable distribution that is purely determined by the transition bias (Eq. (1)). The restoration of the stable phenotype distribution is driven by regrowth of the suppressed phenotypes and therefore occurs considerably slower after growth-independent treatment, i.e., in the direction of a frequency increase of mesenchymal cells (Fig. 5**d**). Such relapses at different rates after distinct treatment types align with our previous findings ^24^ and clinical observations (Section 7 in Chitadze et al. ^40^).

**Figure 5:**
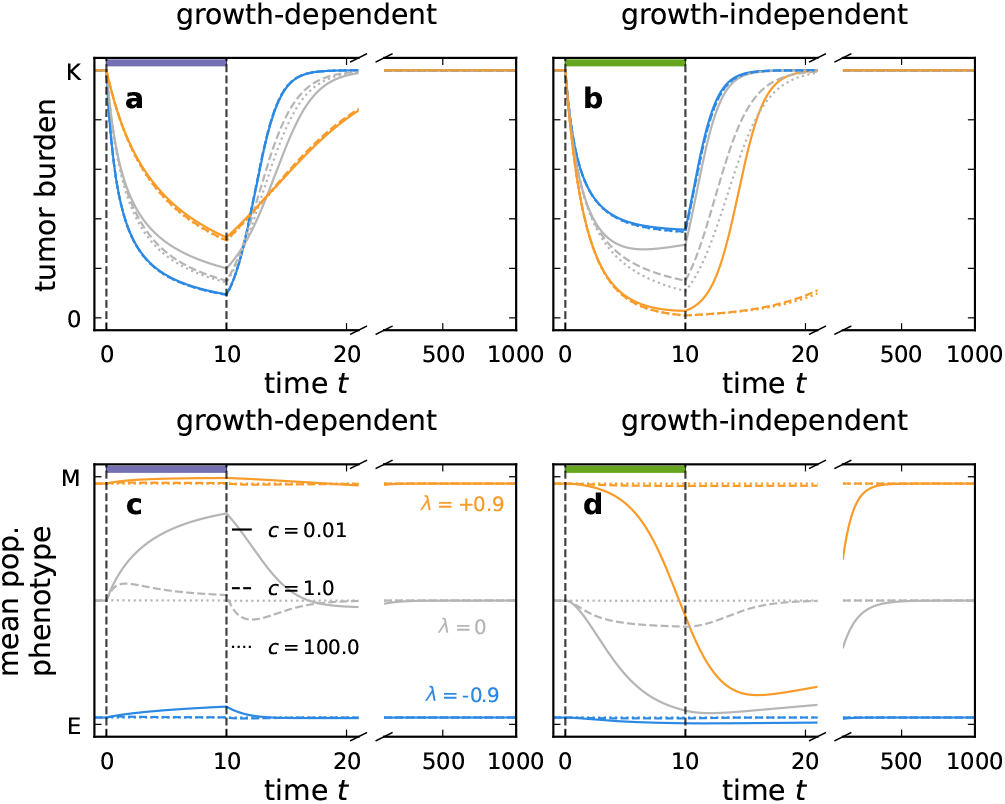
Growth-dependent and growth-independent treatment can transiently alter the phenotype distribution of a tumor. During treatment, the tumor burden reduces (panels **a** and **b)**, and the mean phenotype of the tumor changes if the transition speed *c* is small but restores to the untreated mean phenotype after treatment (panels **c** and **d)**. treatment is applied between *t* = 0 and *t* = 10. Afterward, regrowth is tracked until *t* = 1000. The violet and green horizontal bars indicate growth-dependent and growth-independent treatments. Growth-dependent treatment shifts the mean of the phenotype distribution towards the mesenchymal phenotype as it exerts higher mortality on epithelial cells. Conversely, growth-independent treatment shifts the mean towards the epithelial phenotype as epithelial cells compensate for the mortality by faster growth.

### Tumors with high transition speed are vulnerable to treatment

Exploring a broader range of transition bias and speed, we find that the effect of growth-dependent and growth-independent treatment indeed depends strongly on transition bias and speed (Fig. 6). When a treatment type exerts a particularly high mortality on the most abundant tumor phenotype we refer to the treatment as a phenotype-matched treatment. For instance, growth-dependent treatment exerts the highest mortality on epithelial tumors, *λ* → −1, whereas growth-independent treatment is more phenotype-matched to mesenchymal tumors, *λ* → 1. We find that one-block treatment schemes are most effective, i.e. achieve the strongest reduction in tumor burden, when they are phenotype-matched (Fig. 6, panels **a** and **b**). Applying them for the whole treatment duration thus results in the highest reduction of tumor burden. Higher transition speeds *c* cause a faster replenishment of treatment-sensitive phenotypes from less sensitive phenotypes, entailing a cost of phenotypic plasticity. Thus, treatment efficacy improves for higher transition speeds for single-block treatments.

**Figure 6:**
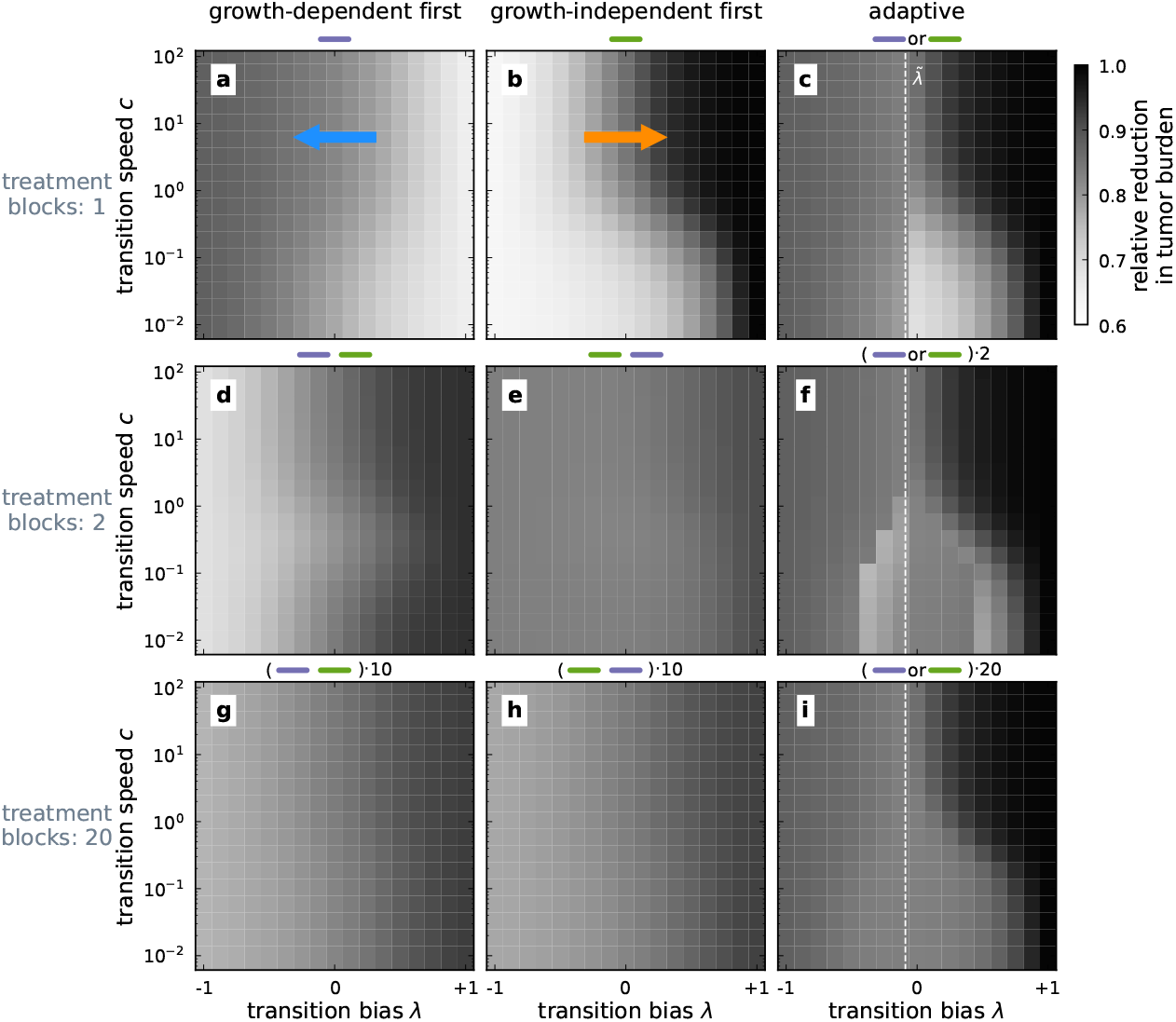
Reduction in tumor burden for different sequential treatment schemes. The reduction in tumor burden relative to the carrying capacity 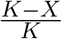 is indicated by the brightness gradient for different combinations of transition bias *λ* and transition speed *c*. Here, *X* is the sum of phenotype abundances at the end of treatment duration (*N* = 3). Darker colors represent a higher reduction and, thus, a better outcome. We evaluate the effect of splitting the treatment period into multiple treatment blocks (rows) and investigate different treatment schemes with either predetined or adaptive treatment sequences (columns). The white dashed line indicates a decision boundary 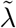 for the adaptive treatment, which is obtained by comparing the mortality of the two treatment types (see text). Adjuvant therapy can alter the phenotype transitions, which in our model translates to changes, for example, to the transition bias *λ*, indicated by the arrows in panels **a** and **b**.

When the phenotype distribution is known, the adaptive treatment scheme can be applied (see Section reatment types and schemes for a detailed description). This treatment scheme selects the treatment type that exerts the higher mortality on the tumor, given the phenotype distribution at the time of treatment choice. We assume that initially, the phenotype distribution has reached its equilibrium given by Eq. (1), which is determined mainly by the transition bias *λ*. For the one-block adaptive treatment scheme only at the beginning of the treatment phase a treatment type has to be determined. Thus, the treatment type decision in the adaptive one-block treatment scheme depends only on the transition bias *λ*. The decision boundary between applying the growth-dependent and the growth-independent treatment in terms of transition bias can be found by comparing the mortality exerted by the two treatment types. The total mortality of growth-dependent treatment 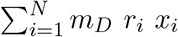 and growth-independent treatment 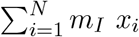 are equal when 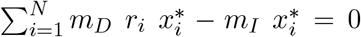. This is a polynomial in the transition bias *λ* with the relevant root 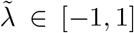 setting the decision boundary between the two treatment types. For our parameters and assumptions on the growth rates of the phenotypes (Table 1), the switch occurs at 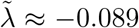. Consequently, in the one-block adaptive treatment (Fig. 6**c**), the reductions of more epithelial tumors with 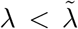 are identical to reductions obtained for the growth-dependent treatment type. More mesenchymal tumors with 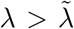 are affected by one-block adaptive treatment identically to the growth-independent treatment.

**Table 1:** Symbols, variables and reference parameters. Deviations from these values are reported where applicable.

| Symbol | Description | Value |
| --- | --- | --- |
| $x_i$ | abundance of $i^{th}$ tumor phenotype | |
| $X$ | sum of all phenotype abundances | $\sum_{i=1}^N x_i$ |
| $N$ | number of tumor phenotypes | 3 |
| $r_i$ | growth rate of $i^{th}$ phenotype | $r_1 - (r_1 - r_N) \frac{i-1}{N-1}$ |
| $r_1$ | growth rate of epithelial cells | 1 |
| $r_N$ | growth rate of mesenchymal cells | $\frac{1}{5}$ |
| $K$ | carrying capacity of a site | 1 |
| $T_{EM}$ | transition rate of $i \rightarrow i + 1$ | $c (1 + \lambda) r_1$ |
| $T_{ME}$ | transition rate of $i \rightarrow i - 1$ | $c (1 - \lambda) r_1$ |
| $c$ | transition speed | 1 |
| $\lambda$ | transition bias | 0 |
| $m_D$ | growth-dependent treatment intensity | 1 |
| $m_I$ | growth-independent treatment intensity | $\approx 0.64$ |

### The effect of sequential multi-block treatment schemes depends on phenotype composition and transition dynamics

To explore the potential of leveraging the treatment-induced phenotype changes, we also investigated sequential treatment patterns that consist of multiple treatment blocks. In the multi-block schemes see (Fig. S2 for tumor burden and mean phenotype dynamics of the two-block scheme), the application duration of the optimal treatment is reduced, thus, resulting in a less effective treatment of tumors with a high transition bias (Fig. 6, panels **d** and **e**). Frequent treatment type alterations result in an intermediate tumor burden reduction across trait space. Although frequent treatment type alteration is not the most effective strategy, it may be desirable when the characteristics of a tumor, such as transition bias *λ* and transition speed *c*, are unknown (Fig. 6, panels **g** and **h**).

For the two-block adaptive treatment, the tumor reduction pattern is more intricate (Fig. 6**f**). At 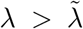, the mortality exerted by growth-independent treatment is higher thus, growth-independent treatment is chosen as the first treatment type. At large transition speed *c* and high transition bias *λ*, growth-independent treatment is also chosen for the second treatment block. Only for a transition bias slightly exceeding 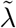 and small *c*, a treatment switch occurs, and the tumor burden reduction is higher than in the single-block scheme. For 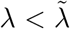, the first treatment type is growth-dependent treatment. Only for small *c* and close to 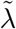 the treatment type is switched after half the treatment period, which interestingly worsens the treatment outcome compared to a single block of only growth-dependent treatment. This choice is possible as the decision criterion for treatment choice is the instantaneous mortality and does not account for the potential future tumor size or the future phenotype distribution, or mortality integrated over time. Since the growth-independent treatment exerts no selection pressure against the proliferative phenotype, switching to this treatment type comes at the risk that soon after the switch, the phenotype distribution will have shifted to a larger epithelial proportion, where the growth-independent treatment is less effective than the growth-dependent treatment. Increasing the number of treatment blocks mediates this risk and achieves the highest tumor burden reduction for small *c* and intermediate *λ* in the adaptive treatment scheme. Different modeling assumptions (unequal competitiveness of phenotypes or phenotype-dependent transition speed) change the effect of treatment only quantitatively (see Sections Phenotype-dependent transition speed, and Unequal competition, Figs. S5 and S8 in the supplementary text).

### The potential of adjuvant treatment

Modifying the process of phenotype transition with adjuvant therapies promises to alter the phenotype distribution of a tumor, thus, rendering the tumor more vulnerable to chemo- or immunotherapy. Transition may be modulated, equivalent of changing transition bias *λ*, by an adjuvant drug that affects the gene regulatory network of epithelial-mesenchymal transitions ^25, 26,29^. (Fig. 6) indicates how modifying the transition bias *λ* with adjuvant drugs can affect the predicted reduction in tumor burden. We find that the tumor burden is reduced more strongly when a single-block growth-dependent treatment is accompanied by an adjuvant drug that makes the tumor more epithelial by decreasing the transition bias *λ* (arrow in Fig. 6**a**). Contrastingly, the effect of the single-block growth-independent treatment can be enhanced by an adjuvant drug that makes the tumor more mesenchymal by increasing the transition bias *λ* (arrow in Fig. 6**b**). For multi-block treatment schemes, we find that increasing transition bias *λ* generally also increases the reduction in tumor burden, albeit this effect depends on the exact pair of transition bias *λ* and speed *c* for the adaptive treatment. Adjuvant therapies that increase the transition speed *c* increase the treatment effect for a single-block treatment, but for two treatment blocks predictions are more difficult as they depend on the treatment sequence and transition bias *λ* and speed *c*.

## 3 Discussion

Cancer cell plasticity increases tumor heterogeneity, facilitates metastasis formation, and often hinders treatment approaches ^41^. We formulated a mathematical model of phenotypic plasticity that motivates a stable heterogeneous phenotype distribution, mainly determined by the relative propensities of phenotype transitions. We found that the heterogeneity generated by this inherent plasticity can explain the transitions between epithelial and mesenchymal phenotypes, which are known to facilitate cancer progression and metastasis ^4,6,13,14,38,42^. Our model provides an alternative hypothesis for the factors driving such transitions, not necessarily relying on the microenvironment yet sensitive to microenvironmental differences. The stability of a heterogeneous phenotype distribution implies that perturbations to this distribution will be reverted. We discuss the consequences of this equilibrium for primary and secondary tumors and their treatment in this section.

### Phenotypic plasticity drives phenotype shifts at the primary and secondary sites of tumor

During metastasis formation, one prominent phenotype transition in cancer cells is the epithelial-mesenchymal transition ^14,23^. Tumors of epithelial origin gain heterogeneity by transitioning to a more mesenchymal phenotype. Selection for motility and invasiveness during the dissemination of cancer cells increases the frequency of the mesenchymal phenotype in the circulating cancer cells. Therefore, the arriving population at the secondary site is mainly constituted of mesenchymal phenotype and gains heterogeneity by recovering more epithelial phenotypes.

Thus, we find that phenotypic plasticity can be responsible for both the epithelial-mesenchymal transition at the primary site and the mesenchymal-epithelial transition at the secondary site. However, microenvironmental differences undoubtedly exacerbate phenotypic changes during the dissemination from primary to secondary site ^8^. Indeed, we can capture much more pronounced phenotype changes by assuming different transition biases *λ*_1_ ≠ *λ*_2_ at primary and secondary sites, resulting in differing stable phenotype distributions.

### Moments of the phenotype distribution guide the choice of phenotype-matched treatment

We investigated growth-dependent and growth-independent treatment types, capturing some mechanisms of chemo- and immunotherapy. We found that the transition bias, and thus the shape of the stable phenotype distribution, determines the most effective treatment type, and sequence of different treatment types. Our model predicts a higher efficiency of growth-dependent treatment for a transition bias towards epithelial phenotypes. Growth-independent treatment becomes more effective for a transition bias towards mesenchymal phenotypes. This connection between treatment effect and the shape of the phenotype distribution can be understood better by considering the moments of the phenotype distribution. Narrow phenotype distributions are well characterized by their mean, which can be used to match treatment type and phenotype.

For example, chemotherapy will be the most effective when there are only proliferative cells. Such a phenotype-matched treatment will perform well for homogeneous tumors in the absence of mismatching terminal phenotypes. To implement this strategy in clinical practice, a thorough characterization of the tumor tissue would be necessary. With the identification of the dominating phenotypes a patient may be stratified for phenotype-matched treatment. Furthermore, patient-derived tumor tissues may be used *in-vitro* to predict the most effective treatment for a particular patient.

However, tumors are often heterogeneous, weakening their treatment response ^43,44^. Interestingly, the third central moment of the phenotype distribution captures the mismatch between a treatment type and the abundance of phenotypes that are not matched by the treatment, as they are far off the mean of the phenotype distribution. When matching the treatment to the mean of the phenotype distribution, an increasing absolute third central moment thus signals the onset of relapse from mismatched phenotypes. In our model, the phenotypic heterogeneity of the stable phenotype distribution is maximal when there is no transition bias, i.e., *λ* = 0. The mismatch between targeted phenotype and unmatched phenotypes, captured by the third central moment, is largest at intermediate transition bias. Accordingly, our model predicts a sweet spot for controlling both heterogeneity and mismatched terminal phenotypes at extreme transition biases *λ* → ±1, where variance and absolute third central moment are simultaneously minimized.

### Molecular drivers of the phenotype transition control the phenotype distribution

Recently, several studies suggested treatment strategies for phenotypically heterogeneous tumors ^24,35,36,45^. These strategies aim to remove or reduce the tumor burden and delay tumor relapse. However, these studies do not target or capitalize on the source of the heterogeneity. The interaction between treatment and phenotype dynamics can potentially be exploited during cancer treatment as the transition bias and, therefore, the stable phenotype distribution can be modulated by an adjuvant therapy. In the context of epithelial-mesenchymal plasticity, the molecular drivers of the epithelial-mesenchymal transition and the mesenchymal-epithelial transitions are known ^17,32^. Altering the concentrations of these drivers in the tumor microenvironment or perturbing the relevant gene regulatory network would thus translate to changing transition bias *λ* and speed *c* which are decisive for the phenotype transition process. Of note, this reasoning applies also to tumor sites that have reached carrying capacity. Although the total abundances of such tumors may stay static, the phenotype distribution can still change ^46^. Also in our model, at carrying capacity, the population growth stops but the transition dynamics can remain in action, capturing the potential turnover in tumor composition. Therefore, it may be possible to regulate tumor phenotypes, reduce metastasis, and achieve a better treatment response by controlling the transition bias with an adjuvant treatment, which, for example, has been shown with TGF-*β* blockage ^25, 26,28^.

### Concentration of molecular drivers may serve as a proxy for phenotype abundances

Building on the decisive effect of such molecules on the phenotype transitions and heterogeneity, their concentration could also be used to characterize tumor heterogeneity. Measuring their concentrations might be practically more feasible than measuring phenotype abundances. For example, in the case of TGF-*β* driven epithelial-mesenchymal transition, the tumor could be characterized by the concentration of TGF-*β* instead of quantifying the number of epithelial, hybrid, and mesenchymal cells. More generally, the state of the relevant gene regulatory network can determine the phenotype distribution, and thus, be used to inform treatment decisions ^29^.

Additionally, if available, the third central moment of the phenotype distribution, i.e., the abundance of mismatched phenotypes, can characterize the relapse. For example, consider a tumor of epithelial cells with a high third central moment. If this tumor is treated with growth-dependent treatment, then the relapse will initially be driven by the growth of mesenchymal cells, eventually, the epithelial cells will take over the population, leading to bi-phasic relapse dynamics ^24,40^. The present model provides a link between the concentration of molecular drivers, equivalent to the transition bias *λ*, and the third central moment of the phenotype distribution.

In this study, we formulated a mathematical model that quantitatively describes the transition between adjacent phenotypes within a population. Phenotype transitions appear in most forms of unicellular organisms. Our findings and the reasoning of targeting phenotype transitions to improve treatment are, hence, applicable beyond cancer, i.e., for pathogen-borne diseases such as bacterial infections. In the current study we make assumptions that allow exposition of our findings, e.g. the phenotypic transitions are not adaptive. Also, we do not model the spread between the primary and secondary cancer sites explicitly. Exchanging some of our simple assumptions for realism, we also discuss the case of unequal competition and growth-dependent transition speed in the supplementary text. The case of unequal competition can be linked to frequency-dependent interactions between tumor phenotypes, which can be captured in the present model by unequal competition coefficients *α*_*i*_. There are compelling studies investigating frequency-dependent interactions ^47–49^ with some quantifying them in patient-derived tumors^50,51^, which could affect treatment outcome and thus, the prognosis of the disease.

To summarize, the present model provides a microenvironment-independent explanation for observed phenotype changes during epithelial-mesenchymal and mesenchymal-epithelial transitions. We highlight the phenotype transitions in a heterogeneous population as a viable clinical target. By targeting the molecular drivers or the relevant gene regulatory network of the phenotype transition, we propose leveraging phenotypic plasticity. Forcing the tumor into a less heterogeneous state, this strategy may improve the treatment efficacy of chemo- and immunotherapies. Supporting the established idea of personalized treatments, our approach thus discusses a practical option to bring the concept into clinical practice.

## 4 Methods

We investigate the onset of metastasis formation mediated by the plasticity between epithelial and mesenchymal phenotypes. We use an ecology-inspired approach to identify tumor phenotypes with species in an ecosystem, drawing on the usefulness of ecological concepts for understanding cancer biology^52,53^. We use the competitive Lotka-Volterra system^54,55^ with additional terms that capture transitions between phenotypes. We go beyond earlier approaches that featured two cell types ^35,36^ and account for the recently established prominence of hybrid phenotypes characterized by both partial epithelial and mesenchymal markers ^15^. This results in a multi-compartment model similar to those presented in Zhou et al. ^33^ and Raatz et al. ^24^, but with added competition and more flexibility in specifying the transition process. Using this model, we examine the epithelial-mesenchymal plasticity and characterize how different properties of the plastic transition shape the resulting tumor heterogeneity.

### Model

We represent the phenotypic heterogeneity as a collection of *N* different tumor phenotypes at different sites of cancer (Eq. (2)). These phenotypes range from the most epithelial phenotype, E, over *N* − 2 hybrid phenotypes H, to the most mesenchymal phenotype, M. We assume that all tumor phenotypes share resources at a given cancer site and that the tumor population follows logistic growth^56^. Additionally, we assume that the tumor cells are phenotypically plastic, i.e., they can change to an adjacent, more epithelial-like, or mesenchymal-like phenotype in bidirectional phenotype transitions.

We track the phenotype abundance *x*_*i*_ over time for each tumor phenotype *i, i* = 1, 2, …, *N*. Each cell of tumor phenotype *i* grows intrinsically at rate *r*_*i*_. The cells switch to the next functionally more mesenchymal phenotype, *i* → *i* + 1, at rate *T*_*EM*_, and to the next functionally more epithelial type, *i* → *i*−1, at rate *T*_*ME*_. The positive transition terms represent additions to a phenotype from adjacent phenotypes. The negative transition terms account for the losses from transitions out of the current phenotype. Resource scarcity in the microenvironment induces a density-dependent, logistic competition captured by 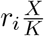, where 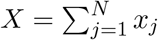 is the total population abundance and *K* is the carrying capacity. We assume that all phenotypes are equally competitive for resources, i.e., the competition term is equal for all phenotypes. Relaxing this assumption does not affect our results qualitatively and is explored in the supplementary text (Section Unequal competition)

Further, we assume linearly decreasing growth rates from epithelial to mesenchymal phenotypes, reflecting the higher proliferation rates of epithelial cells. We scale all growth rates relative to the epithelial growth rate *r*_1_, and express the growth rate for the *i*^*th*^ phenotype as 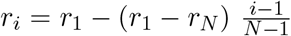. Note that, all the growth rates are equally distributed in the interval [*r*_1_, *r*_*N*_]. The independence of the equilibrium phenotype distribution (Eq. (1)) from the growth rates indicates that the particular choice of the growth rate decline does not affect our results qualitatively.

We assume that the transition rates are equal for all phenotypes and scaled to the epithelial growth rate *r*_1_ with a constant *c*, 0 ≤ *c <* ∞. Thus, transitions faster than proliferation, *c >* 1, can be interpreted as within-generation phenotypic plasticity, e.g. altered patterns of post-transcriptional regulation, and transitions slower than proliferation *c <* 1 can be attributed to transgenerational plasticity, such as epigenetic changes. We explore the effect of growth rate-dependent transition rates in the supplementary text (Section Phenotype-dependent transition speed).

To account for a potential bias in the direction of plastic transitions, we introduce the transition bias *λ*, −1 ≤ *λ* ≤ 1, which determines the probability of a cell switching to the adjacent epithelial or mesenchymal phenotype. A positive transition bias *λ >* 0 implies a bias towards becoming more mesenchymal, *λ <* 0 results in a higher probability of switching to a more epithelial phenotype. With these presumptions, the transition rates become *T*_*EM*_ = *c* (1 + *λ*) *r*_1_ and *T*_*ME*_ = *c* (1 – *λ*) *r*_1_. He parameters of our model are given in Table 1.

### Treatment types and schemes

To investigate the effect of drug treatment on phenotypically plastic tumor cell populations, we apply two different treatment types. The mortality exerted on the cells of a particular phenotype by the two treatment types either does or does not depend on the phenotype’s growth rate. The death rate exerted by growth-dependent treatment is *m*_*D*_ *r*_*i*_, and the death rate by growth-independent treatments is *m*_*I*_. To better compare the phenotype-distribution-dependent treatment effects, we chose *m*_*I*_ such that the reduction of tumor cell abundance at the end of the treatment duration is identical for both treatment types. To find *m*_*I*_, we use *c* = 1, *λ* = 0, *m*_*D*_ = 1, and numerically minimie the difference between the total abundances *X* at the end of treatment duration. Throughout this study, treatment is always applied for a fixed duration of 10 time units.

These assumptions result in a system of ordinary differential equations describing the change in tumor phenotype abundance,

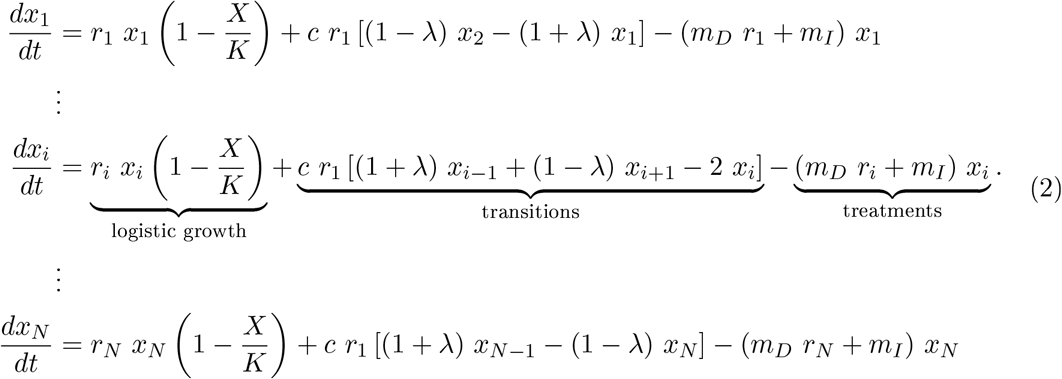

Different drug administration strategies are possible and are of clinical interest. However, in current practice, mostly monotherapies or combined therapies are used, often leading to resistance due to static selection pressures. This motivates us to investigate sequential treatment schemes. The fixed treatment scheme either treats with the same drug throughout the treatment duration or alternates between blocks of different treatment types. The single-block treatment scheme applies either growth-dependent or growth-independent treatment. In the multi-block treatment schemes, we alternate between treatment types and investigate both cases of first treating with growth-dependent or first treating with growth-independent treatment. or the adaptive treatment scheme, at the start of each treatment block, the treatment type is chosen that achieves the higher instantaneous reduction of the whole tumor cell population for the given phenotype distribution ^24^. The cancer cell mortality for growth-dependent treatment is 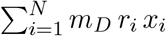 and the mortality exerted by growth-independent treatment is 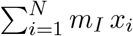. The comparison of these two terms determines the decision for the treatment type.

### Analysis and implementation

We performed stability analysis to find the equilibria of Eqs. (2) and their stability ^57^. He results of this analysis are presented in the Supplementary Text (see Section Stability of the coexistence equilibrium). For *N* = 2 and *N* = 3 stability was contirmed computationally (Fig. S3), for *N >* 3 we show stability using Lyapunov stability analysis. Emporal dynamics of the model (Eqs. (2)) were obtained by numerical integration using the `solve_ivp` function from the `Scipy` library^58^ in Python (version 3.9^59^). Numpy^60^ and Matplotlib ^61^ were used for computation and plotting.

## Data availability

The Figures and dataset generated in this study have been deposited in a zenodo repository with DOI 10.5281/enodo.7989753 ^62^.

## Code availability

The relevant code to generate the Figures and dataset can be found in a zenodo repository with DOI 10.5281/enodo.7989748 ^63^.

## Acknowledgements

All authors acknowledge funding by Deutsche Forschungsgemeinschaft through the Research Training Group “ranslational Evolutionary Research’ (TransEvo) (Project number 400993799, https://gepris.dfg.de/gepris/projekt/400993799) and funding by the Deutsche Krebshilfe (AZ 70112935) to Su.Se. Sa.Sh. is grateful to Dr. Florence Bansept for suggesting alternate parametrization and to Saptarshi Pal for the discussions on stability.

## Author contributions

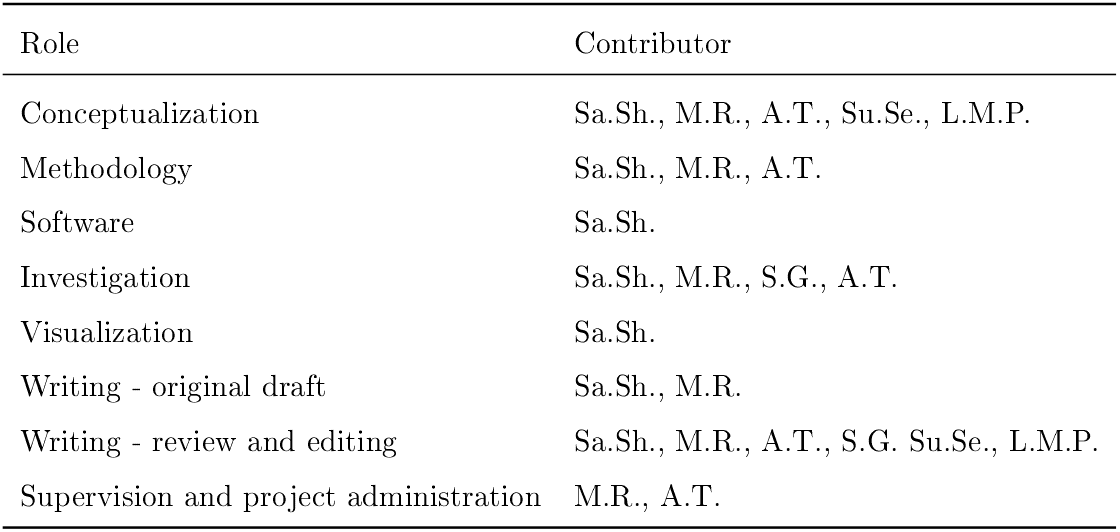

## 5 Supplementary

### 5.1 Supplementary figures

**Figure S1:**
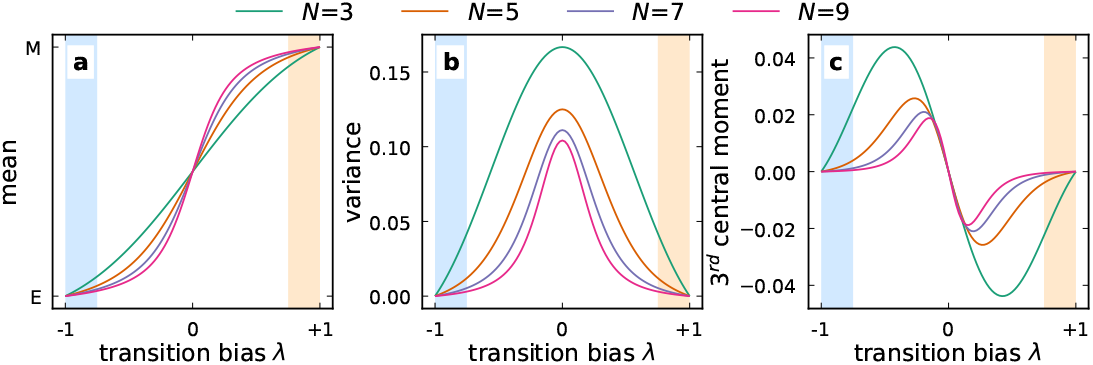
The transition bias affects the phenotypic heterogeneity and abundances of mismatched phenotypes. The stable phenotype distribution 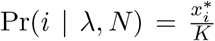, written as a function of transition bias *λ* for different numbers of phenotypes *N*, can be characterized by its moments. Wherever necessary, we scale the phenotype *i* = 0, 1, …, *N* − 1 to a trait axis, 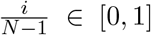, such that 0 always corresponds to epithelial (E) and 1 to mesenchymal M. The first moment of the phenotype distribution is the mean 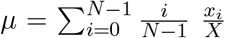. The n^th^ central moment is 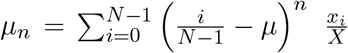. Here, we show the 2^*nd*^ variance and third central moments of the stable phenotype distribution. or measurements, label the phenotype as *i* = 1, 2, …, *N*, measure their abundances *x*_*i*_. Mean is, then, the phenotypes *i* weighted by the measured fractions or relative abundance 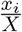. **a** The mean allows the identification of phenotype-matched treatment, e.g., to select growth-dependent treatment for mainly epithelial tumors. **b** The variance represents phenotypic heterogeneity, which often impedes treatment success. **c** The third central moment is a proxy for the abundance of phenotypes far from the mean phenotype that are potentially unmatched by the phenotype-matched treatment. This measure can thus signal when the mean is an inaccurate representation of the phenotype distribution, for example, when it is strongly skewed. Thus, to maximize the efficacy of the phenotype-matched treatment it is favorable to minimize the heterogeneity and inaccuracy of the mean. High transition bias, *λ* → ±1, minimizes both the heterogeneity and the inaccuracy of the mean, indicated by a combination of small variance and a small absolute third central moment. The high transition bias intervals are highlighted with colored background, where the color depicts the type of the most abundant phenotype.

**Figure S2:**
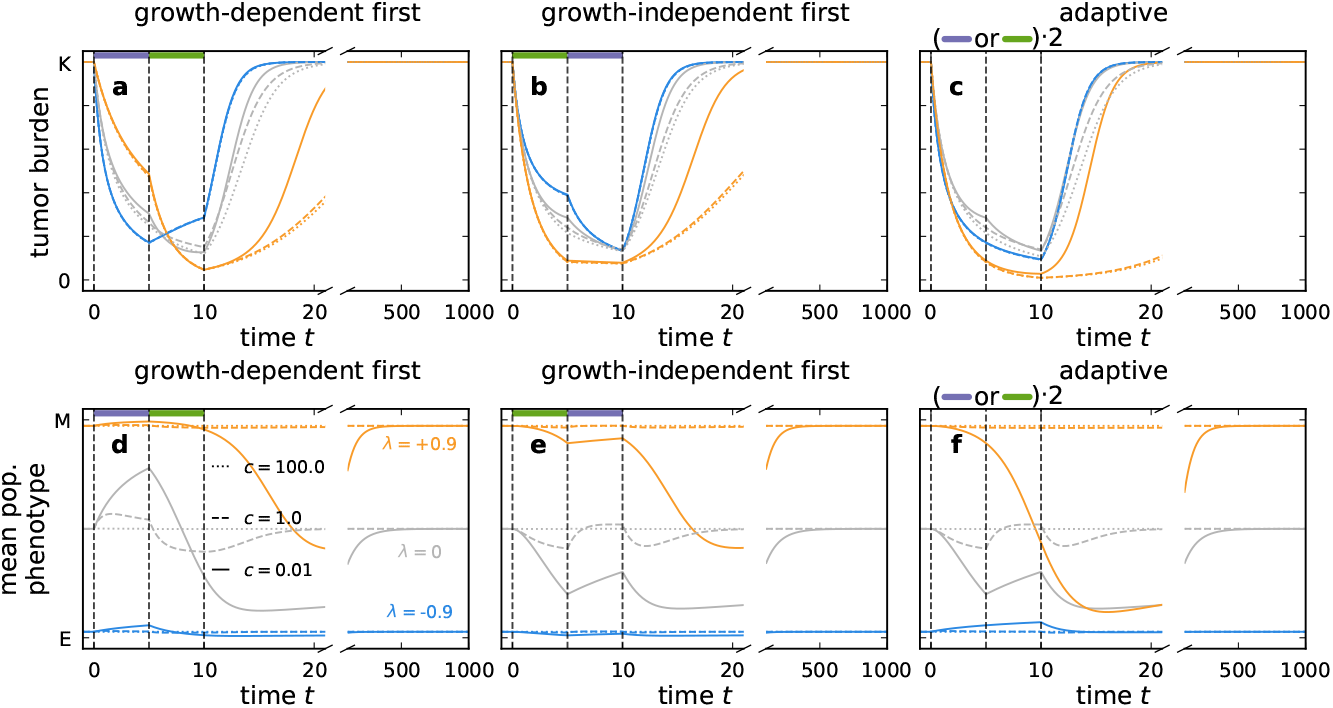
Changing treatment types can increase the efficacy of treatment. The figure shows the effect of sequential two-block treatment on the total tumor abundance (first row) and mean phenotype (second row) during and after treatment for tumors with specific transition bias *λ* (line color) and transition speed *c* (line type). The growth-dependent treatment block is depicted by a violet bar on the top, and the growth-independent block is shown by a green bar. The first block is applied between *t* = 0 and *t* = 5, and the second block is applied between *t* = 5 and *t* = 10. After the treatment duration, the tumor regrowth is tracked until *t* = 1000. or the adaptive treatment scheme, the treatment type is chosen at the beginning of the block based on which type exerts the higher mortality on the present phenotype distribution. In practice, to find the best treatment type, we compare the mortalities exerted by growth-dependent 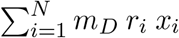, and growth-independent treatment 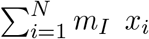. Their difference decides the best treatment type, i.e., 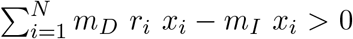 implies that the growth-dependent treatment has higher mortality on the present phenotype distribution. Growth-dependent treatment is chosen for both blocks if the transition bias is low (*λ* = −0.9, blue lines). or unbiased, slow transitions (*λ* = 0 and *c* = 0.01), the treatment type switches from growth-independent to growth-dependent between the first and the second block (solid gray line). The growth-independent treatment type is applied in both treatment blocks for all other parameter combinations. Compared to single-block treatments (Fig. 5), tumor burden reductions at the end of the treatment phase can be higher or lower in two-block treatment schemes, depending on the transition bias and transition speeds, mirroring the complex patterns in Fig. 6. wo-block treatments are thus not always beneficial but can also reduce treatment efficacy if the treatment is not matched with the phenotype distribution.

### 5.2 Stability of the coexistence equilibrium

We investigate the stability of the coexistence phenotype distribution (Eq. (1)) analytically by linear stability analysis for *N* = 2 and *N* = 3 ^57^. For *N >* 3, we compute the eigenvalues using a Mathematica script (`linear_stability.nb`, Supplementary Material) and observe that they are negative for our parameter ranges. For an arbitrary number of phenotypes *N*, we use the Lyapunov stability theory [Chapter 5] ^64^.

#### 5.2.1 Two phenotypes *N* = 2

The following differential equations describe the system with two phenotypes without treatment.

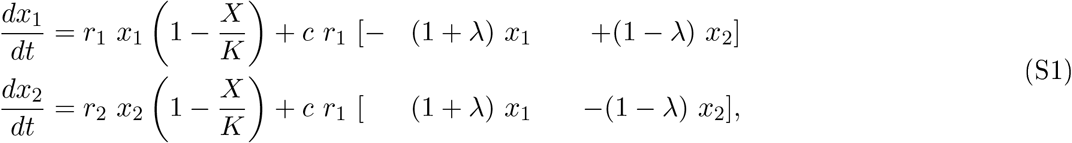

where *X* = *x*_1_ + *x*_2_. The Jacobian **J** of the system at a point (*x*_1_, *x*_2_) is

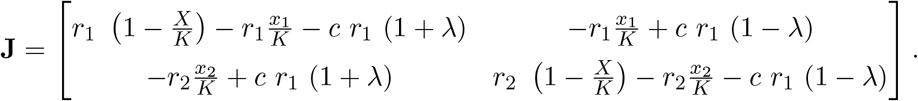

Evaluating the Jacobian at the coexistence equilibrium, *X* = *K*, 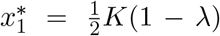, and 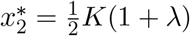 we get

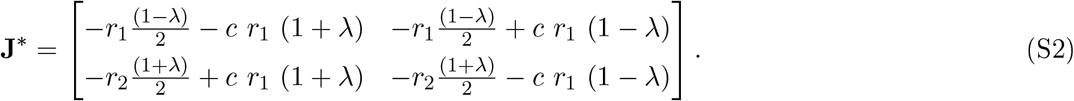

Given that the growth rates are positive by definition *r*_*i*_ *>* 0 and transition bias *λ* ∈ [−1, 1], both the diagonal elements of this matrix are negative, and so must be its trace *τ*. or exposition, using a specific parameter values *r*_1_ = 1, and 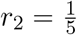, the determinant Δ of the Jacobian at the coexistence fixed point **J**^∗^ is

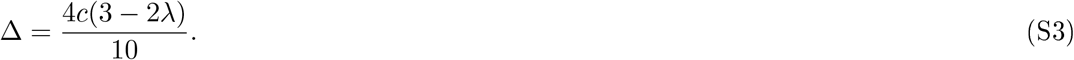

Since transition speed *c* is always positive by definition the determinant Δ of the Jacobian **J**^∗^ is always positive. The stability of this 2D system can be determined from the trace-determinant plane ^57^. The trace is always negative *τ <* 0, and the determinant is always positive Δ *>* 0.

Therefore, the system is linearly stable at the coexistence fixed point.

#### 5.2.2 Three phenotypes *N* = 3

The following differential equations describe the system with three phenotypes where *X* = *x*_1_ + *x*_2_ + *x*_3_.

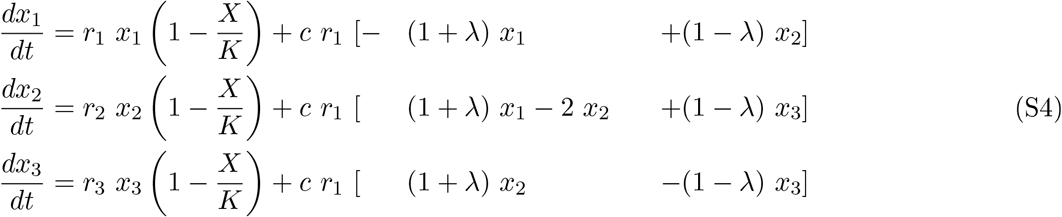

The Jacobian **J** of the system at a point (*x*_1_, *x*_2_, *x*_3_) is

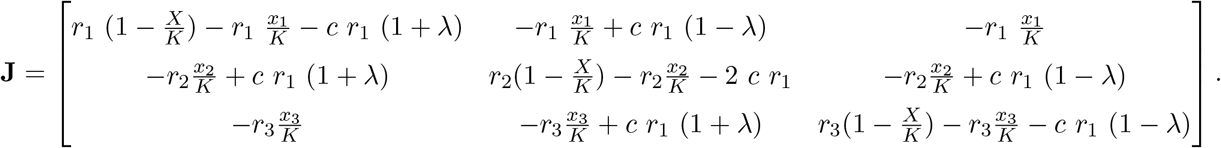

At the coexistence equilibrium, 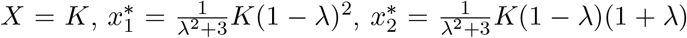, and 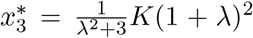, substituting this to get the Jacobian **J**^∗^ at the coexistence fixed point simplities to

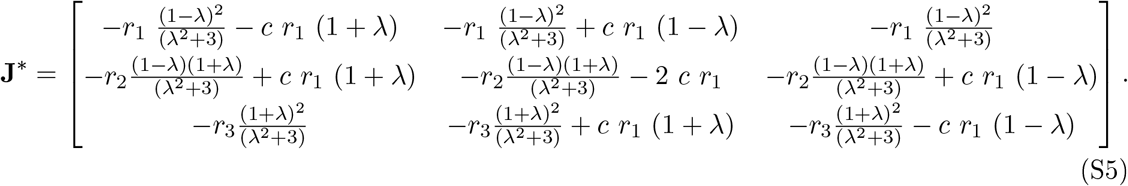

For simplicity, we continue with 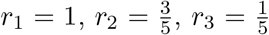, and *c* = 1 to get the characteristic polynomial *P* (*x*). his characteristic polynomial of the Jacobian **J**^∗^ is a 3^*rd*^ order polynomial of the form *P* (*x*) = *x*^3^ + *a*_2_*x*^2^ + *a*_1_*x* + *a*_0_ where

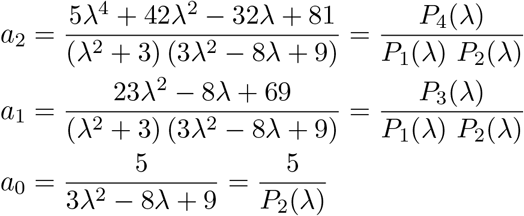

Now, according to the Routh-Hurwit stability criterion ^65^, the system is stable if and only if *a*_2_, *a*_1_, and *a*_0_ are positive and *a*_2_ *a*_1_ − *a*_0_ *>* 0. Above, we have defined four polynomials in *λ*. o show that *a*_2_, *a*_1_, and *a*_0_ are positive, we determine the range of each polynomial for *λ* ∈ [−1, 1]. *P*_1_(*λ*) = *λ*^2^ + 3 is always positive for any real transition bias. The derivative of *P*_2_(*λ*) = 3*λ*^2^–8*λ*+9 is 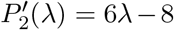 and 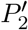 vanishes at 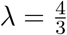 is a parabola opening upwards, 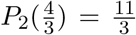 is the minimum, and *P*_2_(*λ*) is positive for any real transition bias. The derivative of *P*_3_(*λ*) = 23*λ*^2^ − 8*λ* + 69 is 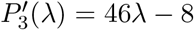 and 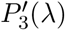 vanishes at 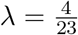. As also *P*_3_(*λ*) is a parabola opening upwards, 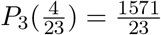 is a minimum, and also *P*_3_(*λ*) is positive for a real transition bias.

**Figure S3:**
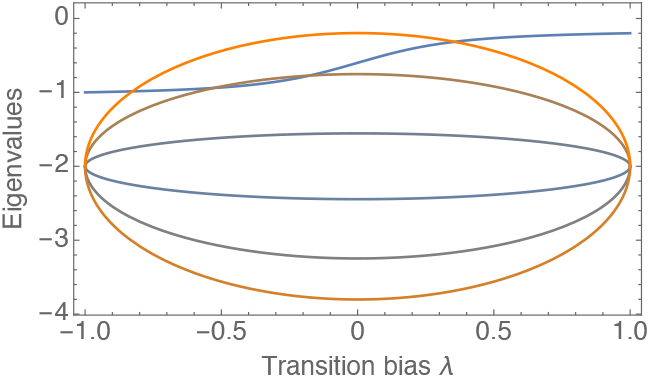
The coexistence phenotype distribution is locally stable for any number of phenotypes. The Jacobian matrix of the system Eq. (2) is evaluated at the coexistence phenotype distribution, 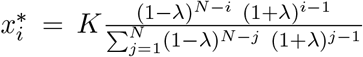, with *r* = 1, *c* = 1. or the number of phenotypes *N* = 7, all 7 eigenvalues of the Jacobian matrix at the coexistence phenotype distribution are negative. Thus, the coexistence phenotype distribution is locally stable.

The first derivative of *P*_4_(*λ*) = 5*λ*^4^ + 42*λ*^2^ − 32*λ* + 81 is 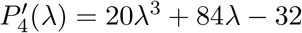 and has a unique real root *λ*^∗^ ≈ 0.37. The second derivative of *P*_4_(*λ*) is 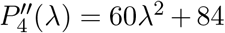 which is always positive for a real transition bias *λ*. Since 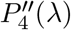 has no roots there is no inflection and *P*_4_(*λ*) is always concave up. Thus, *P*_4_(*λ*^∗^) ≈ 75 is the minimum as the curvature is always positive, and *P*_4_(*λ*) is positive for a real transition bias.

Therefore, *a*_2_, *a*_1_, and *a*_0_ are positive for a transition bias −1 *< λ <* 1. inally, we need *a*_2_ *a*_1_ − *a*_0_ *>* 0 to show that all eigenvalues of the Jacobian **J**^∗^ have a negative real part.

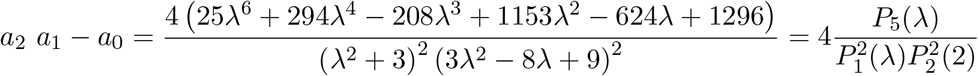

Following a similar procedure as above reveals that *P*_5_(*λ*) is always concave up and positive for any real transition bias. Thus, all Routh-Hurwit conditions are satified, and the coexistence equilibrium is a linearly stable fixed point.

#### 5.2.3 N > 3 phenotypes

We provide a Mathematica ^66^ script that computes the eigenvalues for phenotypes up to order of *N* = 10 and plots them for the range of the transition bias *λ* ∈ [−1, 1]. As an example, Fig. S3 shows the eigenvalues for *N* = 7, which are all negative for realistic ranges of our model parameters. The script is available at DOI: 10.5281/enodo.7294563. For a more rigorous argument we There provide a Lyapunov stability analysis. The system in Eq. 2 can be written more compactly in matrix form as

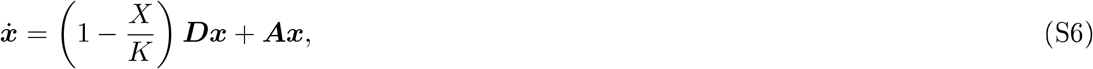

where

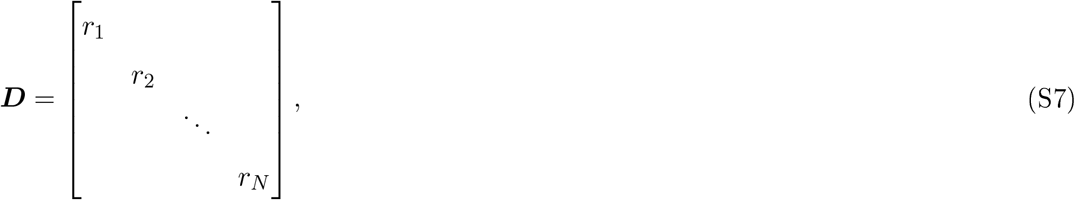

and

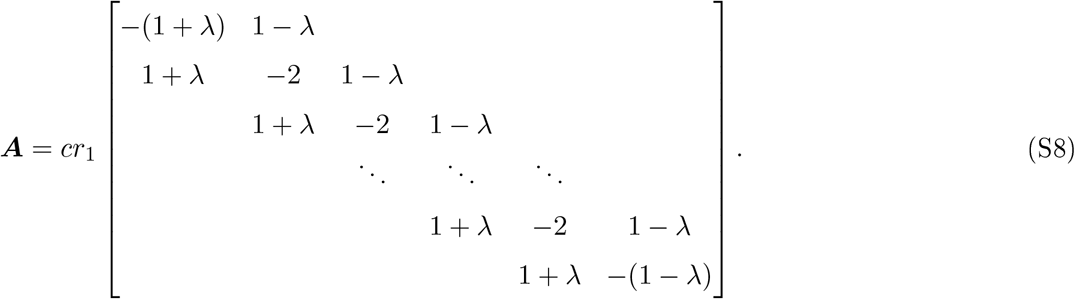

The matrix

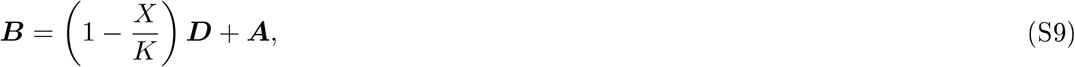

allows us to write the system in Eq. S6 as 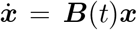. Since all off-diagonal entries of ***B***(*t*) are non-negative for *t* ≥ 0, the system in Eq. S6 is positive Theorem 3.1 ^67^ and ***x***(*t*) ≥ 0 and ***x***(*t*) ≠ 0 for all *t* ≥ 0 meaning that, when the system starts with non-negative initial conditions, it stays non-negative throughout.

To study the dynamics of this system, define the scalar function *V* as

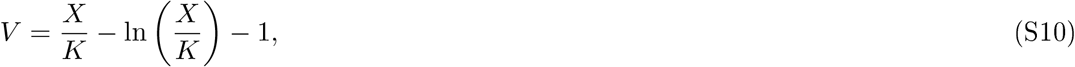

and note that *V >* 0 for *X* ≠ *K* while *V* = 0 for *X* = *K*. Differentiating,

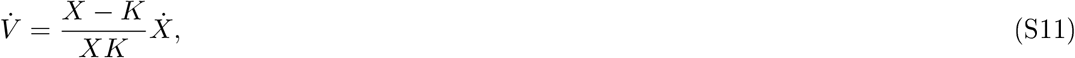

where from Eq. 2, the definition of *X*, and telescoping,

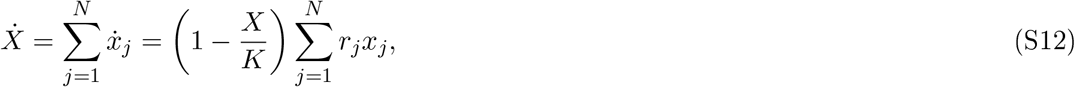

so that we can write the right-hand side of Eq. S11 in closed form as

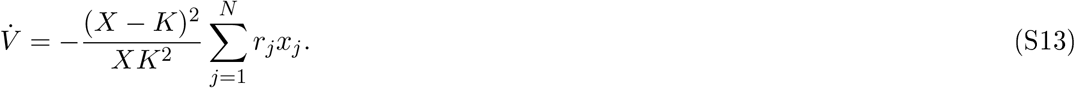

This expression shows that the derivative of *V* is negative whenever *X* ≠ *K*, while it vanishes at *X* = *K*. Therefore, *V* is a Lyapunov function for the dynamics of *X* and *X*^∗^ = *K* is a globally stable equilibrium of *X*. Notably, this stability result is independent of the dynamics of the system that originate from the second term on the right-hand side of Eq. S6.

Due to this result, the system in Eq. S6 eventually approaches the linear system

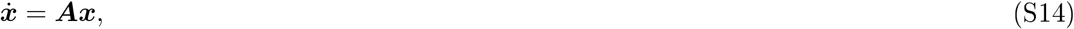

as *t* → ∞, where ***A*** is time independent. The system of Eq. (2) has fixed points when the two terms, growth dynamics and transition dynamics, in Eq. (S6) vanish simultaneously. All phenotype contiguration satisfying *X* = *K* make the first term vanish.

The transition matrix ***A*** can be seen as an irreducible generator of a continuous-time Markov chain. For *λ* ∈ (−1, +1), the irreducibility follows from the fact that you can go from any phenotype *i* to any other phenotype *j* ≠ *i* through some path on the directed graph of ***A***. Then, by Lemma A.2 from 68, 0 is an eigenvalue of ***A*** with multiplicity 1, and all other eigenvalues have negative real parts. For *λ* ± 1, the multiplicity 1 of eigenvalue 0 follows from the diagonal of the resulting triangular matrix. Multiplying any vector that is parallel to the unique eigenvector ***v***, corresponding to the 0 eigenvalue, to the right side of ***A*** will yield 0. Therefore Eq. (1), obtained by scaling eigenvector ***v*** such that the components of ***v*** sum to *K*, is the unique solution to Eq. (S6) for all *N*.

**Figure S4:**
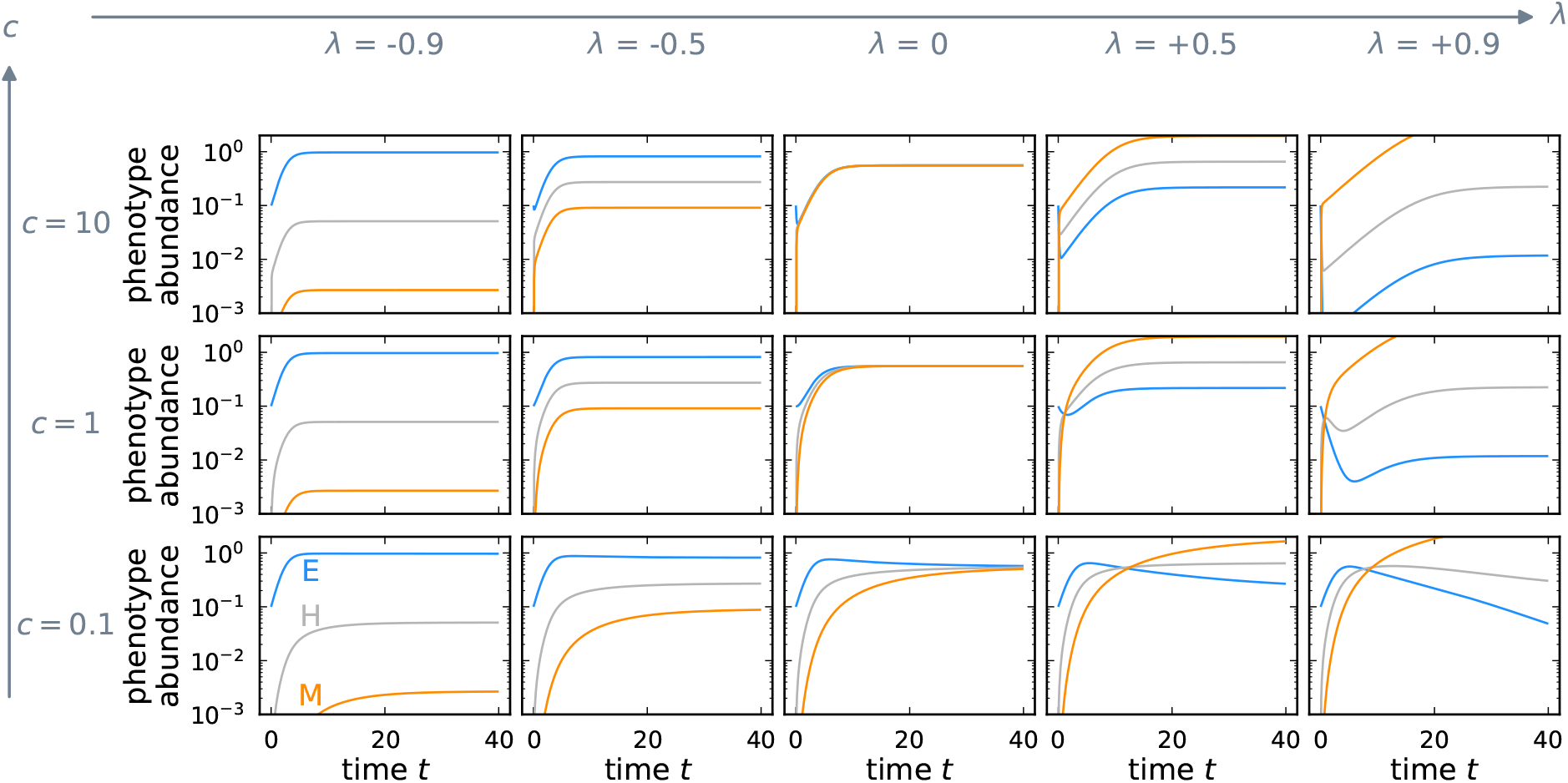
Unequal competition allows tumor burdens higher than the carrying capacity. The panels show approach to the stable phenotype distribution from the same initial condition,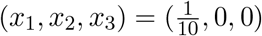, for different combinations of transition speed *c* and transition bias *λ*. We assume that cells consume resources proportional to their growth rate, *α*_*i*_ = *r*_*i*_ · (1 *time unit*). Since mesenchymal cells consume fewer resources, the higher the mesenchymal proportion, the more cells can be sustained above the carrying capacity.

### 5.3 Model extensions

#### 5.3.1 Unequal competition

In the original model, we assumed that all phenotypes share the resources identically. However, resource utilization of phenotypes may differ and lead to a different competitiveness of phenotypes. To account for this aspect, There we extend the model by including phenotype-specific competition coefficients *α*_*i*_. Thus, we replace *X* in Eqs. (2) with 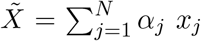 and obtain for the untreated dynamics

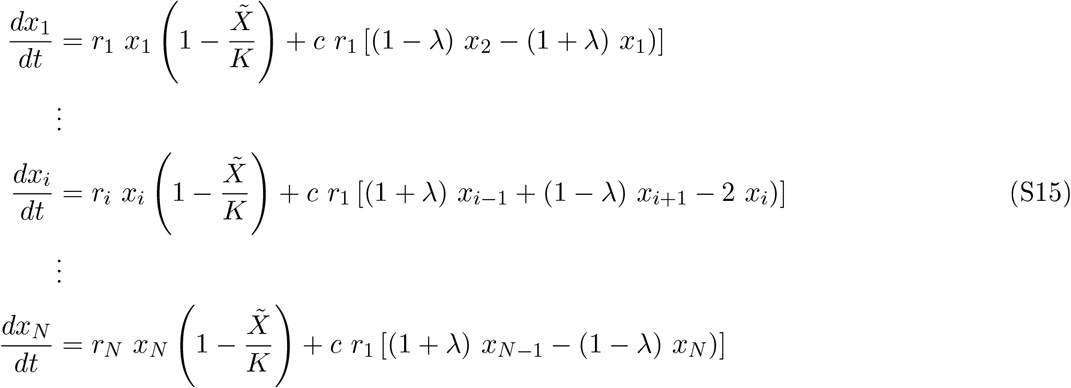

This set of equations has a saddle point and a stable equilibrium similar to the original model. The state with no cells is an unstable equilibrium. The stable equilibrium is given by 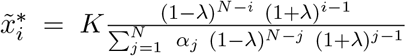. Note that the stable equilibrium is obtained without any assumptions on the competition coefficients and therefore is valid for any non-negative competition coefficients. The competition coefficients *α*_*i*_ affect only the carrying capacity in the form of a constant scaling factor 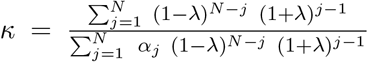. The stable phenotype distribution with unequal competition is, thus, the equilibrium abundances 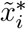 divided by this scaled carrying capacity *κK*, e.g. 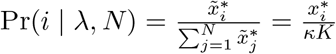. Remarkably, the transition bias *λ* solely determines the stable phenotype distribution also in the case of unequal competition between phenotypes.

**Figure S5:**
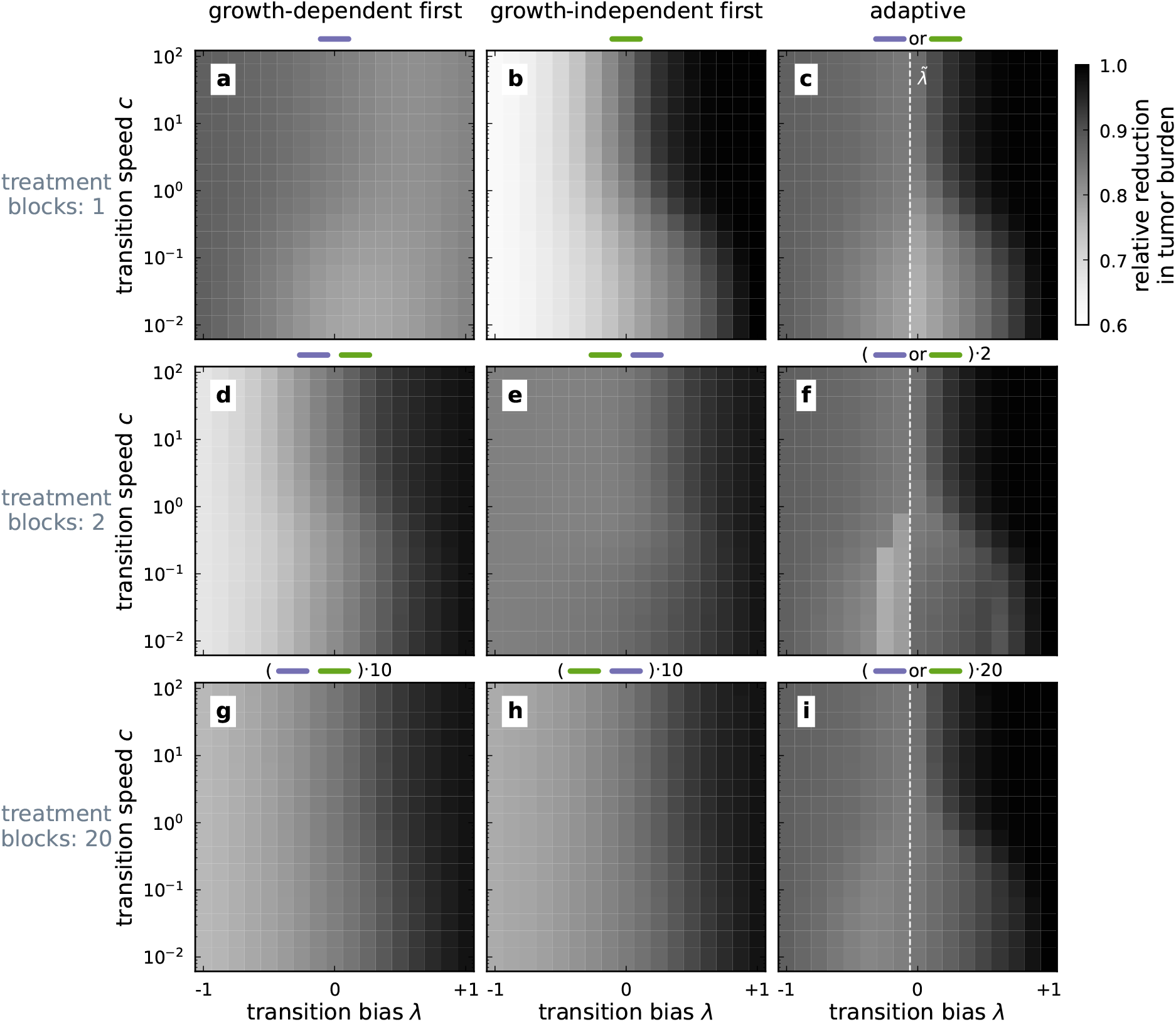
Unequal competition improves treatment efficiencies for unbiased and mesenchymal-biased tumors. The heatmap shows the relative reduction in tumor burden from the abundance fixed point 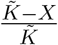 at the stable phenotype distribution for unequal competition where 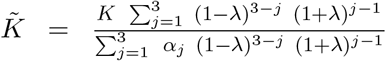. The intensity of growth-independent treatment is *m*_*I*_ ≈ 0.65. Darker colors represent a higher reduction and, thus, a better outcome. We evaluate the effect of splitting the treatment period into multiple treatment blocks (rows) and investigate different treatment schemes with either predetined or adaptive treatment sequences (columns).

**Figure S6:**
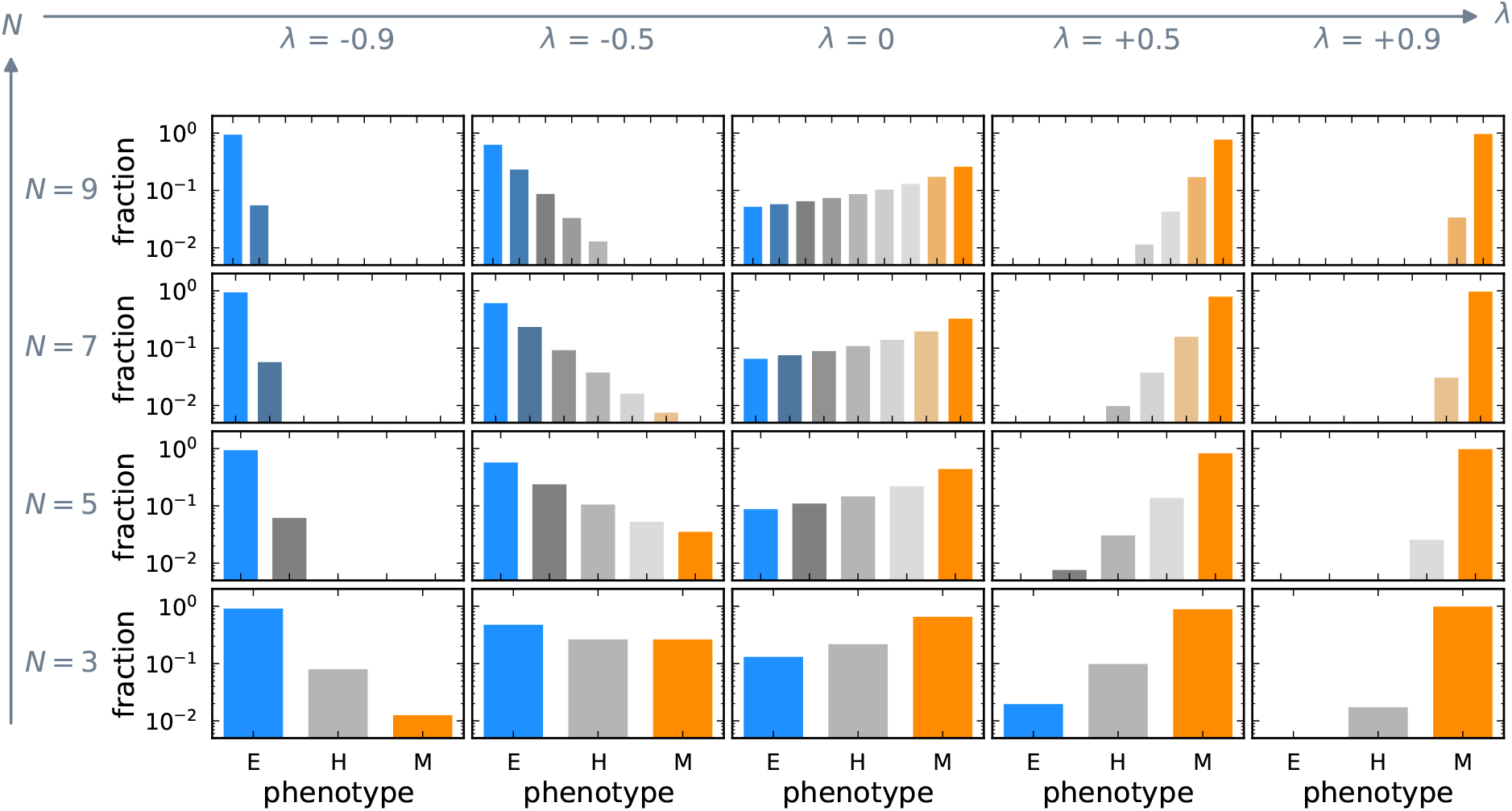
Phenotype-dependent transition speed does not qualitatively affect the stable phenotype distribution. Each panel shows the equilibrium distribution of phenotypes for a fixed value of the transition bias and the number of phenotypes. The stable phenotype distribution changes with the transition bias *λ*. When there is no transition bias to switch to either an epithelial or a mesenchymal-like phenotype, i.e., *λ* = 0, the stable distribution is not uniform as faster proliferating cells transition faster. Transition bias towards the epithelial-like phenotype, *λ <* 0, leads to an increase in epithelial cells. Conversely, a transition bias towards mesenchymal-like phenotypes, *λ >* 0, leads to an increase in mesenchymal cells.

Notably, now the abundances, particularly of the mesenchymal phenotype, can exceed the carrying capacity *K*, i.e. *κ >* 1, up to an effective carrying capacity given by 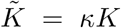 (Fig. S4). Consider the case where mesenchymal-like cells use fewer resources than epithelial-like cells. Namely, we assume that the competition coefficients are proportional to the growth rate. Rescaling the competition coefficients then gives *α*_*i*_ = *r*_*i*_ η 1 (time unit) for the *i*^*th*^ phenotype. Note that the units of growth rate and competition coefficients are different requiring the multiplication by 1 (time unit). This assumption incorporates a classical *r* − *K* trade-o into our model with more proliferative but less competitive epithelial cells and less proliferative but more competitive mesenchymal cells ^69^. This higher abundance of less proliferative but more competitive mesenchymal phenotypes increases the treatment efficiencies for intermediate and high transition biases (Fig. S5). Similarly, when a phenotype is harmful to others *α*_*j*_ *>* 1 the equilibrium abundance can be lower than the carrying capacity *κ <* 1.

#### 5.3.2 Phenotype-dependent transition speed

Here, we investigate the effect of using *c r*_*i*_ for the transition speed of phenotype *i* instead of *c r*_1_, i.e., phenotypic transitions are coupled to cell divisions, and the transition rates are scaled by the growth rate of each phenotype.

**Figure S7:**
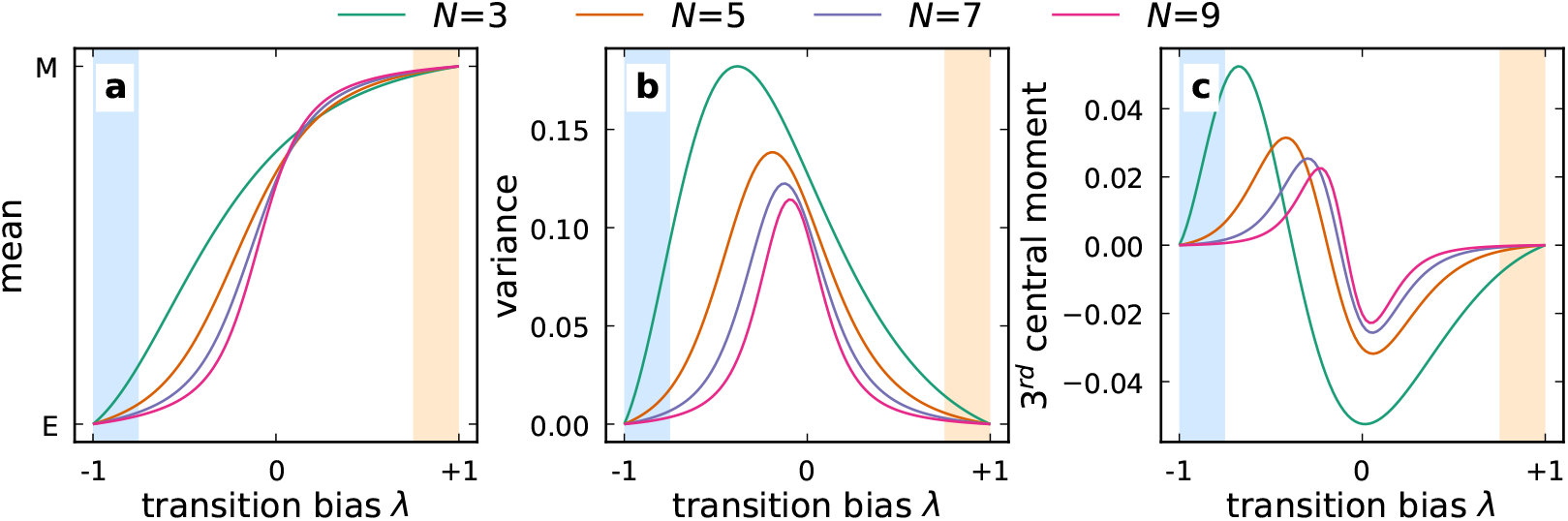
Phenotype-dependent transition speed decreases the abundance of epithelial phenotypes. Plot specifics are identical to Fig. S1. Including phenotype-dependent transition speed breaks the symmetry of the stable phenotype distribution. The overall mesenchymal proportion increases as fast-proliferating epithelial cells transition faster into mesenchymal cells. Thus, the growth-dependent treatment type loses efficiency due to a worse phenotype match for a tumor with phenotype-dependent transition speed.

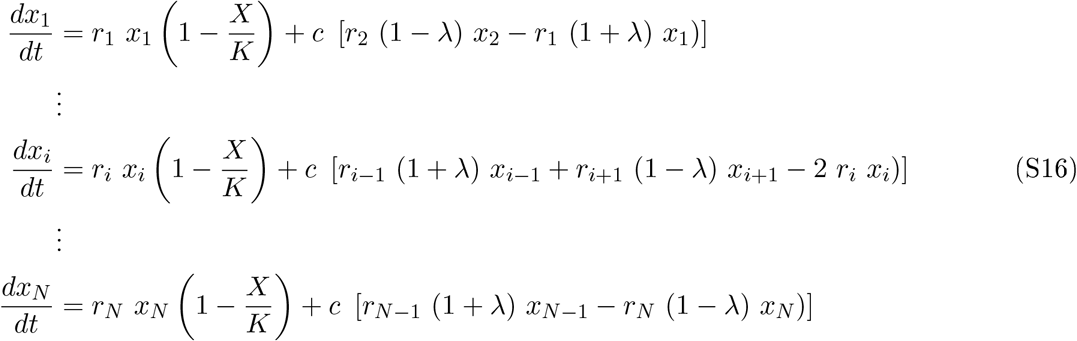

This set of equations has two equilibria similar to the original model. One equilibrium is a saddle-node equilibrium of extinction, and the other is a stable equilibrium with a fixed phenotype distribution. This stable equilibrium is given by 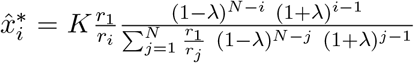. The model dynamics with phenotype-dependent transition speed are qualitatively similar to the original model. In this case, however, the stable phenotype distribution depends on the phenotype growth rate, increasing the fraction of more mesenchymal phenotypes (Fig. S6). This leads to stable phenotype distributions with negative third central moment for a larger transition bias range (Fig. S7) and an efficiency loss of purely growth-dependent treatment (Fig. S8).

**Figure S8:**
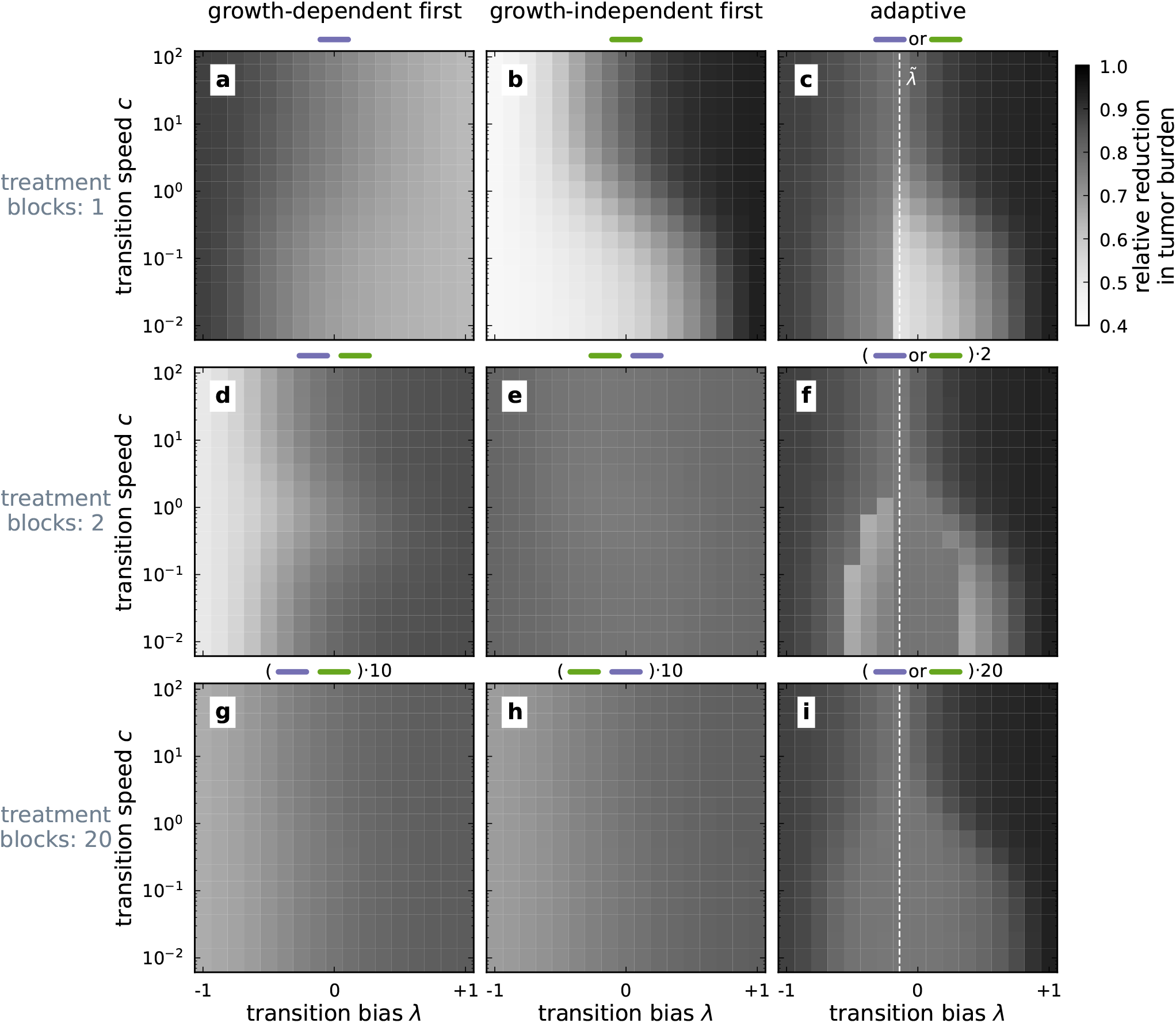
Phenotype-dependent transition speed improves the treatment efficiency. The heatmap shows the relative reduction in tumor burden from the abundance fixed point 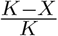 at the stable phenotype distribution for phenotype-dependent transition speed. The intensity of growth-independent treatment is *m*_*I*_ ≈ 0.50. Darker colors represent a higher reduction and, thus, a better outcome. We evaluate the effect of splitting the treatment period into multiple treatment blocks (rows) and investigate different treatment schemes with either predefined or adaptive treatment sequences (columns). The negative effect of transition speed is amplified by phenotype-dependent transition speed. Thus, the overall growth rate of the tumor is decreased, further reducing the tumor burden at the end of the treatment duration.

## References

[1] Hanna Dillekås, Michael S. Rogers, and Oddbjørn Straume. Are 90D of deaths from cancer caused by metastases & Cancer Medicine, 8(12):5574–5576, September 2019. ISSN 2045-7634, 2045-7634. doi: 10.1002/cam4.2474.

[2] Rebecca L. Siegel, 8imberly D. Miller, and Ahmedin Jemal. Cancer statistics, 2020. C Cancer Journal or Clinicians, 70(1):7–30, January 2020. ISSN 0007-9235, 1542-4863. doi: 10.3322/caac.21590.

[3] Hyuna Sung, Jacques Ferlay, Rebecca L. Siegel, Mathieu Laversanne, Isabelle Soerjomataram, Ahmedin Jemal, and Freddie Bray. Global Cancer Statistics 2020: GLOBOCAN Estimates of Incidence and Mortality Worldwide for 36 Cancers in 185 Countries. C Cancer Journal or Clinicians, 71(3):209–249, May 2021. ISSN 0007-9235, 1542-4863. doi: 10.3322/caac.21660.

[4] Gaorav P. Gupta and Joan Massagué. Cancer Metastasis: Building a Framework. Cell, 127(4):679–695, November 2006. ISSN 00928674. doi: 10.1016/j.cell.2006.11.001.

[5] Robert A. Weinberg. the Biology of Cancer. Garland Science, Taylor F Francis Group, New York, second edition edition, 2014. ISBN 978-0-8153-4219-9 978-0-8153-4220-5.

[6] S. Turajlic and C. Swanton. Metastasis as an evolutionary process. Science, 352(6282):169–175, April 2016. ISSN 0036-8075, 1095-9203. doi: 10.1126/science.aaf2784.

[7] YelyHaveta Shlyakhtina, 8atherine L. Moran, and Maximiliano M. Portal. Genetic and Non-Genetic Mechanisms Underlying Cancer Evolution. Cancers, 13(6):1380, March 2021. ISSN 2072-6694. doi: 10.3390/cancers13061380.

[8] Group Young Researchers in Inflammatory Carcinogenesis, Anna Maxi Wandmacher, Anne-Sophie Mehdorn, and Susanne Sebens. The Heterogeneity of the Tumor Microenvironment as Essential Determinant of Development, Progression and Therapy Response of Pancreatic Cancer. Cancers, 13(19):4932, September 2021. ISSN 2072-6694. doi: 10.3390/cancers13194932.

[9] Francisco Caiado, Bruno Silva-Santos, and Håkan Norell. Intra-tumour heterogeneity - going beyond genetics. e FbJJS Journal, 283(12):2245–2258, June 2016. ISSN 1742464X. doi: 10.1111/febs.13705.

[10] M J West-Eberhard. Phenotypic Plasticity and the Origins of Diversity. Annual Review of Ecology and Systematics, 20(1):249–278, November 1989. ISSN 0066-4162. doi: 10.1146/annurev.es.20.110189.001341.

[11] Beat B. Fischer, Marek Kwiatkowski, Martin Ackermann, Jasmin Lrismer, Severin RoMer, Marc J. F. Suter, Rik L. Eggen, and Blake Matthews. Phenotypic plasticity influences the eco-evolutionary dynamics of a predator-prey system. bcolo, 95(11):3080–3092, November 2014. ISSN 0012-9658. doi: 10.1890/14-0116.1.

[12] Alf Giese, Melinda A. Loo, Nhan Tran, Dorothy Haskett, Stephen W. Coons, and Michael E. Berens. Dichotomy of astrocytoma migration and proliferation. International Journal of Cancer, 67(2):275–282, July 1996. ISSN 00207136, 10970215. doi: 10.1002/(SICI)1097-0215(19960717)67:2<275::AID-IJC20>3.0.C02-9.

[13] Xiaofeng Zheng, Julienne L. Carstens, Jiha Kim, Matthew Scheible, Judith Kaye, Hikaru Sugimoto, Chia-Chin Wu, Valerie S. LeBleu, and Raghu Kalluri. Epithelial-to-mesenchymal transition is dispensable for metastasis but induces chemoresistance in pancreatic cancer. Nature, 527(7579):525–530, November 2015. ISSN 0028-0836, 1476-4687. doi: 10.1038/nature16064.

[14] Mohit Kumar Jolly, Kathryn E. Ware, Shivee Gilja, Jason A. Somarelli, and Herbert Levine. EMT and MET: Necessary or permissive for metastasis? Molecular Oncology, 11(7):755–769, July 2017. ISSN 1574-7891, 1878-0261. doi: 10.1002/1878-0261.12083.

[15] Mohit Kumar Jolly, Sendurai A. Mani, and Herbert Levine. Hybrid epithelial/mesenchymal phenotype(s): The EfittestG for metastasis? Biochimica et Biophysica Acta (BBA) - Reviews on Cancer, 1870(2):151–157, December 2018. ISSN 0304419X. doi: 10.1016/j.bbcan.2018.07.001.

[16] Deli Hong, Andrew J. Fritz, Sayyed K. ;aidi, Andre J. Wijnen, JeJrey A. Nickerson, Anthony N. Imbalzano, Jane B. Lian, Janet L. Stein, and Gary S. Stein. Epithelial-to-mesenchymal transition and cancer stem cells contribute to breast cancer heterogeneity. Journal of Cellular Physiology, 233(12):9136–9144, December 2018. ISSN 0021-9541, 1097-4652. doi: 10.1002/jcp.26847.

[17] Anushka Dongre and Robert A. Weinberg. New insights into the mechanisms of epithelial-mesenchymal transition and implications for cancer. Nature Reviews Molecular Cell Biology, 20(2):69–84, February 2019. ISSN 1471-0072, 1471-0080. doi: 10.1038/s41580-018-0080-4.

[18] Nicole Vincent Jordan, Gary L. Johnson, and Amy N. Abell. Tracking the intermediate stages of epithelial-mesenchymal transition in epithelial stem cells and cancer. Cell Cycle, 10(17):2865–2873, September 2011. ISSN 1538-4101, 1551-4005. doi: 10.4161/cc.10.17.17188.

[19] R Y-J Huang, M K Wong, T Z Tan, K T Kuay, A H C Ng, V Y Chung, Y-S Chu, N Matsumura, H-C Lai, Y F Lee, W-J Sim, C Chai, E Pietschmann, S Mori, J J H Low, M Choolani, and J P Thiery. An EMT spectrum defines an anoikis-resistant and spheroidogenic intermediate mesenchymal state that is sensitive to e-cadherin restoration by a src-kinase inhibitor, saracatinib (A;D0530). Cell Death & Disease, 4(11):e915–e915, November 2013. ISSN 2041-4889. doi: 10.1038/cddis.2013.442.

[20] Mohit Kumar Jolly, Jason A. Somarelli, Maya Sheth, Adrian Biddle, Satyendra C. Tripathi, Andrew J. Armstrong, Samir M. Hanash, Sharmila A. Bapat, Annapoorni Rangarajan, and Herbert Levine. Hybrid epithelial/mesenchymal phenotypes promote metastasis and therapy resistance across carcinomas. Pharmacology E0 Therapeutics, 194:161-184, February 2019. ISSN 01637258. doi: 10.1016/j.pharmthera.2018.09.007.

[21] Gianluca Selvaggio, Sara Canato, Archana Pawar, Pedro T. Monteiro, Patrícia S. Guerreiro, M. Manuela Brás, Florence Janody, and Claudine Chaouiya. Hybrid Epithelial-Mesenchymal Phenotypes Are Controlled by Microenvironmental Factors. Cancer Research, 80(11):2407–2420, June 2020. ISSN 0008-5472, 1538-7445. doi: 10.1158/0008-5472.CAN-19-3147.

[22] Hanah Goetz, Juan R. Melendez-Alvarez, Luonan Chen, and Xiao-Jun Tian. A plausible accelerating function of intermediate states in cancer metastasis. PLOS Computational Biology, 16(3):e1007682, March 2020. ISSN 1553-7358. doi: 10.1371/journal.pcbi.1007682.

[23] M. Angela Nieto, Ruby Yun-Ju Huang, Rebecca A. Jackson, and Jean Paul Thiery. EMT: 2016. Cell, 166(1):21–45, June 2016. ISSN 00928674. doi: 10.1016/j.cell.2016.06.028.

[24] Michael Raatz, Saumil Shah, Guranda Chitadze, Monika Brüggemann, and Arne Traulsen. The impact of phenotypic heterogeneity of tumour cells on treatment and relapse dynamics. PLOS Computational Biology, 17(2):e1008702, February 2021. ISSN 1553-7358. doi: 10.1371/journal.pcbi.1008702.

[25] JiePiong Liu, Shan Liao, Benjamin Diop-Frimpong, Wei Chen, Shom Goel, Kamila NaQerova, Marek Ancukiewicz, Yves Boucher, Rakesh K. Jain, and Lei Xu. TGF-β blockade improves the distribution and efficacy of therapeutics in breast carcinoma by normalizing the tumor stroma. Proceedings of the National Academy of Sciences, 109(41): 16618-16623, decmber 2012. ISSN 0027-8424, 1091-6490. doi: 10.1073/pnas.1117610109.

[26] Michael Pickup, Sergey Novitskiy, and Harold L. Moses. The roles of TGFβ in the tumour microenvironment. Nature Reviews Cancer, 13(11):788–799, November 2013. ISSN 1474-175X, 1474-1768. doi: 10.1038/nrc3603.

[27] Weikang Wang, Dante Poe, Yaxuan Yang, Thomas Hyatt, and Jianhua Xing. Epithelial-to-mesenchymal transition proceeds through directional destabilization of multidimensional attractor. eLife, 11:e74866, February 2022. ISSN 2050-084X. doi: 10.7554/eLife.74866.

[28] Hyang-Mi Lee, Hye-Jin Lee, and Ji-Eun Chang. Inflammatory Cytokine: An Attractive Target for Cancer Treatment. Biomedicines, 10(9):2116, August 2022. ISSN 2227-9059. doi: 10.3390/biomedicines10092116.

[29] Kishore Hari, Varun Ullanat, Archana Balasubramanian, Aditi Gopalan, and Mohit Kumar Jolly. Landscape of epithelial-mesenchymal plasticity as an emergent property of coordinated teams in regulatory networks. eLife, 11: e76535, October 2022. ISSN 2050-084X. doi: 10.7554/eLife.76535.

[30] Mingyang Lu, Mohit Kumar Jolly, Herbert Levine, JosE N. Onuchic, and Eshel Ben-Jacob. MicroRNA-based regulation of epithelial-hybrid-mesenchymal fate determination. Proceedings of the National Academy of Sciences, 110(45): 18144–18149, November 2013. ISSN 0027-8424, 1091-6490. doi: 10.1073/pnas.1318192110.

[31] Abhijeet P. Deshmukh, Suhas V. Vasaikar, Katarzyna Tomczak, Shubham Tripathi, Petra Den Hollander, Emre Arslan, Priyanka Chakraborty, Rama Soundararajan, Mohit Kumar Jolly, Kunal Rai, Herbert Levine, and Sendurai A. Mani. Identification of EMT signaling cross-talk and gene regulatory networks by single-cell RNA sequencing. Proceedings of the National Academy of Sciences, 118(19):e2102050118, May 2021. ISSN 0027-8424, 1091-6490. doi: 10.1073/pnas.2102050118.

[32] Ayalur Raghu Subbalakshmi, Sarthak Sahoo, Isabelle McMullen, Aaditya Narayan Saxena, Sudhanva Kalasapura Venugopal, Jason A. Somarelli, and Mohit Kumar Jolly. KLF4 Induces Mesenchymal-Epithelial Transition (MET) by Suppressing Multiple EMT-Inducing Transcription Factors. Cancers, 13(20):5135, October 2021. ISSN 2072-6694. doi: 10.3390/cancers13205135.

[33] Da Zhou, Yue Luo, David Dingli, and Arne Traulsen. The invasion of de-differentiating cancer cells into hierarchical tissues. PLOS Computational Biology, 15(7):e1007167, July 2019. ISSN 1553-7358. doi: 10.1371/journal.pcbi.1007167.

[34] Piyush B. Gupta, Christine M. Fillmore, Guozhi Jiang, Sagi D. Shapira, Kai Tao, Charlotte Kuperwasser, and Eric S. Lander. Stochastic State Transitions Give Rise to Phenotypic Equilibrium in Populations of Cancer Cells. Cell, 146 (4):633–644, August 2011. ISSN 00928674. doi: 10.1016/j.cell.2011.07.026.

[35] Xin Li and D. Thirumalai. A mathematical model for phenotypic heterogeneity in breast cancer with implications for therapeutic strategies. Journal of The Royal Society interface, 19(186):20210803, January 2022. ISSN 1742-5662. doi: 10.1098/rsif.2021.0803.

[36] Tyler Cassidy, Daniel Nichol, Mark Robertson-Tessi, Morgan Craig, and Alexander R. A. Anderson. The role of memory in non-genetic inheritance and its impact on cancer treatment resistance. PLOS Computational Biology, 17 (8):e1009348, August 2021. ISSN 1553-7358. doi: 10.1371/journal.pcbi.1009348.

[37] Bo Ma, Alan Wells, and Amanda M. Clark. The pan-therapeutic resistance of disseminated tumor cells: Role of phenotypic plasticity and the metastatic microenvironment. Seminars in Cancer Biology, 60:138–147, February 2020. ISSN 1044579X. doi: 10.1016/j.semcancer.2019.07.021.

[38] Mark Esposito, Shridar Ganesan, and Yibin Kang. Emerging strategies for treating metastasis. Nature Cancer, 2(3): 258–270, March 2021. ISSN 2662-1347. doi: 10.1038/s43018-021-00181-0.

[39] Linnea C Franssen and Mark A J Chaplain. A mathematical multi-organ model for bidirectional epithelial-mesenchymal transitions in the metastatic spread of cancer. IMA Journal of Applied athematics, 85 (5):724–761, September 2020. ISSN 0272-4960, 1464-3634. doi: 10.1093/imamat/hxaa022.

[40] Guranda Chitadze, Anna Laqua, Marcus Lettau, Claudia D Baldus, and Monika BrLggemann. Bispecific antibodies in acute lymphoblastic leukemia therapy. Hxpert Review of Hematology, 13(11):1211–1233, November 2020. ISSN 1747-4086, 1747-4094. doi: 10.1080/17474086.2020.1831380.

[41] Saniya Deshmukh and Supreet Saini. Phenotypic Heterogeneity in Tumor Progression, and Its Possible Role in the Onset of Cancer. Drontiers in & enetics, 11:604528, November 2020. ISSN 1664-8021. doi: 10.3389/fgene.2020.604528.

[42] Jawad Fares, Mohamad Y. Fares, Hussein H. Khachfe, Hamza A. Salhab, and Youssef Fares. Molecular principles of metastasis: A hallmark of cancer revisited. Signal Transduction and Targeted Therapy, 5(1):28, December 2020. ISSN 2059-3635. doi: 10.1038/s41392-020-0134-x.

[43] Alissa Greenbaum, David R. Martin, Thèrése Bocklage, Ji-Hyun Lee, Scott A. Ness, and Ashwani Rajput. Tumor Heterogeneity as a Predictor of Response to Neoadjuvant Chemotherapy in Locally Advanced Rectal Cancer. Clinical Colorectal Cancer, 18(2):102–109, June 2019. ISSN 15330028. doi: 10.1016/j.clcc.2019.02.003.

[44] Kerry-Ann McDonald, Tsutomu Kawaguchi, Qianya Qi, Xuan Peng, Mariko Asaoka, Jessica Young, Mateusz Opyrchal, Li Yan, Santosh Patnaik, Eigo Otsuji, and Kazuaki Takabe. Tumor Heterogeneity Correlates with Less Immune Response and Worse Survival in Breast Cancer Patients. Annals of Surgical Oncology, 26(7):2191–2199, July 2019. ISSN 1068-9265, 1534-4681. doi: 10.1245/s10434-019-07338-3.

[45] Yannick Viossat and Robert Noble. A theoretical analysis of tumour containment. Nature Ecology & Evolution, 5(6): 826–835, June 2021. ISSN 2397-334X. doi: 10.1038/s41559-021-01428-w.

[46] Michael Raatz and Arne Traulsen. Promoting extinction or minimizing growth? The impact of treatment on trait trajectories in evolving populations. Evolution, page qpad042, March 2023. ISSN 0014-3820, 1558-5646. doi: 10.1093/evolut/qpad042.

[47] I.P.M. Tomlinson. Game-theory models of interactions between tumour cells. b’uropean Journal of Cancer, 33(9): 1495–1500, August 1997. ISSN 09598049. doi: 10.1016/S0959-8049(97)00170-6.

[48] Ipm Tomlinson and Wf Bodmer. Modelling the consequences of interactions between tumour cells. Hritish Journal of Cancer, 75(2):157–160, January 1997. ISSN 0007-0920, 1532-1827. doi: 10.1038/bjc.1997.26.

[49] Marvin A. Böttcher, Janka Held-Feindt, Michael Synowitz, Ralph Lucius, Arne Traulsen, and Kirsten Hattermann. Modeling treatment-dependent glioma growth including a dormant tumor cell subpopulation. HMC Cancer, 18(1): 376, December 2018. ISSN 1471-2407. doi: 10.1186/s12885-018-4281-1.

[50] Audrey R. Freischel, Mehdi Damaghi, Jessica J. Cunningham, Arig Ibrahim-Hashim, Robert J. Gillies, Robert A. Gatenby, and Joel S. Brown. Frequency-dependent interactions determine outcome of competition between two breast cancer cell lines. Scientific Reports, 11(1):4908, March 2021. ISSN 2045-2322. doi: 10.1038/s41598-021-84406-3.

[51] Artem Kaznatcheev, Jeffrey Peacock, David Basanta, Andriy Marusyk, and Jacob G. Scott. Fibroblasts and alectinib switch the evolutionary games played by non-small cell lung cancer. Nature b’cology f3 Evolution, 3(3):450–456, February 2019. ISSN 2397-334X. doi: 10.1038/s41559-018-0768-z.

[52] David Basanta and Alexander R. A. Anderson. Exploiting ecological principles to better understand cancer progression and treatment. Iinterface ocus, 3(4):20130020, August 2013. ISSN 2042-8898, 2042-8901. doi: 10.1098/rsfs.2013.0020.

[53] Kirill S. Korolev, Joao B. Xavier, and Jeff Gore. Turning ecology and evolution against cancer. Nature Reviews Cancer, 14(5):371–380, May 2014. ISSN 1474-175X, 1474-1768. doi: 10.1038/nrc3712.

[54] S. Smale. On the differential equations of species in competition. Journal of Mathematical Hiology, 3(1):5–7, 1976. ISSN 0303-6812, 1432-1416. doi: 10.1007/BF00307854.

[55] Josef Hofbauer and Karl Sigmund. Evolutionary Games and Population Dynamics. Cambridge Lniversity Press, Mrst edition, May 1998. ISBN 978-0-521-62365-0 978-0-521-62570-8 978-1-139-17317-9. doi: 10.1017/CBO9781139173179.

[56] Vinay G. Vaidya and Frank J. Alexandro. Evaluation of some mathematical models for tumor growth. international Journal of Hio-Medical Computing, 13(1):19–35, January 1982. ISSN 00207101. doi: 10.1016/0020-7101(82)90048-4.

[57] Steven H. Strogatz. Nonlinear Dynamics and Chaos. CRC Press, zeroth edition, May 2018. ISBN 978-0-429-96111-3. doi: 10.1201/9780429492563.

[58] Pauli Virtanen, Ralf Gommers, Travis E. Oliphant, Matt Haberland, Tyler Reddy, David Cournapeau, Evgeni Burovski, Pearu Peterson, Warren Weckesser, Jonathan Bright, Stéfan J. van der Walt, Matthew Brett, Joshua Wilson,K. Jarrod Millman, Nikolay Mayorov, Andrew R. J. Nelson, Eric Jones, Robert Kern, Eric Larson, C J Carey, ilhan Polat, Yu Feng, Eric W. Moore, Jake VanderPlas, Denis Laxalde, Josef Perktold, Robert Cimrman, Ian Henriksen, E. A. Quintero, Charles R. Harris, Anne M. Archibald, Antônio H. Ribeiro, Fabian Pedregosa, Paul van Mulbregt, SciPy 1.0 Contributors, Aditya Vijaykumar, Alessandro Pietro Bardelli, Alex Rothberg, Andreas Hilboll, Andreas Kloeckner, Anthony Scopatz, Antony Lee, Ariel Rokem, C. Nathan Woods, Chad Fulton, Charles Masson, Christian Häggström, Clark Fitzgerald, David A. Nicholson, David R. Hagen, Dmitrii V. Pasechnik, Emanuele Olivetti, Eric Martin, Eric Wieser, Fabrice Silva, Felix Lenders, Florian Wilhelm, G. Young, Gavin A. Price, Gert-Ludwig Ingold, Gregory E. Allen, Gregory R. Lee, Hervé Audren, Irvin Probst, Jörg P. Dietrich, Jacob Silterra, James T Webber, Janko Slavič, Joel Nothman, Johannes Buchner, Johannes Kulick, Johannes L. Schönberger, José Vinícius de Miranda Cardoso, Joscha Reimer, Joseph Harrington, Juan Luis Cano Rodríguez, Juan Nunez-Iglesias, Justin Kuczynski, Kevin Tritz, Martin Thoma, Matthew Newville, Matthias Kümmerer, Maximilian Bolingbroke, Michael Tartre, Mikhail Pak, Nathaniel J. Smith, Nikolai Nowaczyk, Nikolay Shebanov, Oleksandr Pavlyk, Per A. Brodtkorb, Perry Lee, Robert T. McGibbon, Roman Feldbauer, Sam Lewis, Sam Tygier, Scott Sievert, Sebastiano Vigna, Stefan Peterson, Surhud More, Tadeusz Pudlik, Takuya Oshima, Thomas J. Pingel, Thomas P. Robitaille, Thomas Spura, Thouis R. Jones, Tim Cera, Tim Leslie, Tiziano Zito, Tom Krauss, Utkarsh Upadhyay, Yaroslav O. Halchenko, and Yoshiki Vázquez-Baeza. SciPy 1.0: Fundamental algorithms for scientific computing in Python. Nature Methods, 17(3):261–272, March 2020. ISSN 1548-7091, 1548-7105. doi: 10.1038/s41592-019-0686-2.

[59] Guido van Rossum and Fred L. Drake. The Python Language Reference. Number Pt. 2 in Python Documentation Manual / Guido van Rossum; Fred L. Drake [Ed.]. Python Software Foundation, Hampton, NH, release 3.0.1 [repr.] edition, 2010. ISBN 978-1-4414-1269-0.

[60] Charles R. Harris, K. Jarrod Millman, Stefan J. van der Walt, Ralf Gommers, Pauli Virtanen, David Cournapeau, Eric Wieser, Julian Taylor, Sebastian Berg, Nathaniel J. Smith, Robert Kern, Matti Picus, Stephan Hoyer, Marten H. van Kerkwijk, Matthew Brett, Allan Haldane, Jaime Fernandez del Rio, Mark Wiebe, Pearu Peterson, Pierre Gerard- Marchant, Kevin Sheppard, Tyler Reddy, Warren Weckesser, Hameer Abbasi, Christoph Gohlke, and Travis E. Oliphant. Array programming with NumPy. Nature, 585(7825):357–362, September 2020. ISSN 0028-0836, 1476-4687. doi: 10.1038/s41586-020-2649-2.

[61] John D. Hunter. Matplotlib: A 2D Graphics Environment. Computin in Science & Engineerin, 9(3):90–95, 2007. ISSN 1521-9615. doi: 10.1109/MCSE.2007.55.

[62] Saumil Shah, Lisa-Marie Philipp, Stefano Giaimo, Susanne Sebens, Arne Traulsen, and Michael Raatz. Understanding and leveraging phenotypic plasticity during metastasis formation - Dataset, May 2022.

[63] Saumil Shah, Lisa-Marie Philipp, Stefano Giaimo, Susanne Sebens, Arne Traulsen, and Michael Raatz. Understanding and leveraging phenotypic plasticity during metastasis formation - Code. Zenodo, May 2023.

[64] Shankar Sastry. Nonlinear Systems, volume 10 of Interdisciplinary Applied Mathematics. Springer New York, New York, NY, 1999. ISBN 978-1-4419-3132-0 978-1-4757-3108-8. doi: 10.1007/978-1-4757-3108-8.

[65] Feliks R. Gantmacher. The Theory of Matrices. Vol. 2, volume 2. American Mathematical Soc, Providence, RI, reprinted edition, 2009. ISBN 978-0-8218-2664-5.

[66] Inc. Wolfram Research. Mathematica. Wolfram Research, Inc., 2021.

[67] D.S. Bernstein and S.P. Bhat. Nonnegativity, reducibility, and semistability of mass action kinetics. In Proceedings of the 38th IEEE Conference on Decision and Control (Cat. No.99CH36304), volume 3, pages 2206-2211, Phoenix, AZ, USA, 1999. IEEE. ISBN 978-0-7803-5250-6. doi: 10.1109/CDC.1999.831248.

[68] George Yin and Qing Zhang. Continuous Time Markov Chains and Applications: A Two-Time-Scale Approach. Number 37 in Applications of Mathematics. Springer, New York Berlin Heidelberg, 2nd ed edition, 2013. ISBN 978-1-4899-9118-8 978-1-4614-4345-2.

[69] Eric R. Pianka. On r- and K-Selection. The American Naturalist, 104(940):592–597, 1970. ISSN 00030147, 15375323.

